# Elucidating the short and long-term mechanical response of the cell nucleus with a hybrid-viscoelastic model

**DOI:** 10.1101/542274

**Authors:** Daniel Pérez-Calixto, Erika González-Villa, Edgar Jiménez-Díaz, Nathalia Serna-Márquez, Genaro Vázquez-Victorio, Mathieu Hautefeuille

## Abstract

The mechanical properties of the nucleus play an important role in all the processes of a cell and impact greatly its decisions, functions and phenotype. It is then important to understand how internal and external stresses can modify them. To study the mechanical response of the nucleus at different timescales, a hybrid viscoelastic model integrating both continuum mechanics and soft glass matter theory is developed. It indeed accounts for the instantaneous viscoelastic response of the structural components of the nucleus as well as the active response of the nuclear envelope and the dynamic reorganization of the cytoskeleton at different timescales. This model can describe adequately the nuclear deformation caused by substrate stiffness in primary hepatocytes and HepG2 cells in culture up to 5 days. It also reveals that the increase of nuclear strain in the long term implies nuclear softening (a phenomenon intensified on stiffer substrates), simultaneously with an increase of the dissipative properties of the nucleus, offering stability. Finally, in the context of soft glassy theory, the model suggests that processes of aging and mechanical memory of the cell may be originated by the dissipative capacity of the nuclei.

## Introduction

Adherent cells perceive and respond constantly to mechanical strain and stresses from their microenvironment through various mechanisms that enable the conversion of those stimuli to biochemical signals^1–4^. Moreover, it has been demonstrated that cells not only sense but also generate stresses that are responsible for their internal organization^5–7^. This process of balancing internal and external stresses commonly results in a mechanically stable state^8^, originated by the constant dynamic reorganization of the cytoskeleton with a purpose to maintain cellular shape and function^7,9–11^. In particular, the existence of a direct mechanical link via the adhesion complex and cytoskeleton has been shown between the cell microenvironment and its nucleus: thanks to this chain the external mechanical signals are hence transmitted to and perceived by the nucleus^12,13^. Because its role is to store, organize and protect the genetic material of the cell^14,15^, any alteration of the mechanical properties of the nucleus would not only affect the integrity of the genome^14–20^, hence the dysregulation of gene expression, but also some alterations of its structural components are associated with diverse pathologies like certain forms of cancer^21–23^, multiple laminopathies^18,24,25^ and cardiac diseases^26,27^. Today, we know that the mechanical properties of the cell nucleus are mainly dictated by its nuclear envelope (NE) and the chromatin acetylation state (Ch) ^15,28–31^. The NE is principally composed of lamins A/C and lamins B1 and B2, that not only provide structural stability to the nucleus but also play an important role in the regulation of different processes such as: cellular differentiation, genome organization, replication of genes and embryonic development^19,32–36^. Recent evidence has shown that a cascade of biochemical pathways depending on contractility may modify the structure of chromatin, hence its mechanical characteristics by way of the translocation of epigenetic factors to the nucleus^30,37,38^. Thanks to the development of different types of mechanical characterization techniques^39^, it has been possible to study the rheology of cell nucleus and the spatial architecture of cell nucleus^29^. However, several challenges remain unsolved to fully understand, describe and model the mechanical behavior of the cellular nucleus in its natural form, i.e. inside a cell. Indeed, considering that most of the existing techniques used to determine the mechanical characteristics of a material must deform it or impose a direct stress upon its surface^40^, the techniques using a physical contact with a very small cell nucleus that is naturally protected by a surrounding network of interconnected cytoskeleton fibers^12,29,41^ may overestimate or underestimate the measured values, introduce artefacts or even alter the system under test and its context^42^. For example, the measured Young’s modulus values obtained for isolated nuclei differ, in some cases, up to 50% when compared with nuclei measured inside the cells^43^. For this reason, non invasive measurement techniques have been recently developed to improve the characterization, such as Brillouin microscopy and magnetic resonance elastography, aiming at establishing an empirical correlation between physiological and pathological processes. Because these dynamic techniques do not allow to obtain the Young’s modulus directly but rather measure the longitudinal modulus, a continuous calibration of the systems and a correct interpretation of resulting data are necessary^40^.

Given that several studies have shown that the cell nucleus can present both a reversible^44,45^ and irreversible^46,47^ mechanical behavior, the object of the present study is to elucidate the long-term evolution of the nuclear shape due to physiological and pathological mechanical conditions. This is performed exploring the impact of the extracellular matrix (ECM) stiffness on the nuclear strain dynamics, since modifications of the ECM stiffness is related to pathological events ^48–50^. In particular, the physiological condition is emulated by freshly isolated rat Wistar hepatocytes seeded directly on a soft (1 kPa) polyacrylamide hydrogel, and the pathological condition was represented by using a stiffer hydrogel (23 kPa). This experimental setup approaches in a simple manner the healthy and fibrotic conditions of a liver tissue ^51,52^. Additionally, to disclose if the nuclear response to these specific conditions is dependent of the cell mechanical pre-state, the same assay is performed using the HepG2 cell line, which comes from a human hepatocellular carcinoma and has been maintained and expanded always on a stiff polystyrene culture plates (~2 GPa)^53^. In particular, the possible manifestation of an irreversible mechanical behavior due to the nuclear strain dynamics or a previous mechanical configuration (an established mechanical preconfiguration defined by stiffness and morphology in a previous, original state) was studied in the long term, using a hybrid viscoelastic model (H-VM). The model incorporates the reported viscoelastic nature of NE and Ch^54–58^ with relaxation times in the range of minutes^55^. Also, the active behavior of nuclear lamins was considered and integrated, using a gene circuit based on the one that was proposed by Dingal and Discher^59^. In order to compare the classical and macroscopic time-independent approach with the mesoscopic and time-dependent description of the nuclear rheology, the long-term nucleus mechanical behavior (in the range of hours and days) was included in the H-VM in two different ways, in order to represent each view. First, a description of an hyperelastic material is used; a theory described by parameters which are scalar functions of the deformation in thermal equilibrium^60–65^. This theory model originates from the observations of the mechanical behavior of matter at a bulk-level and its constitutive equations may be deduced from a thermodynamic potential. However, the macroscopic nature of hyperelastic materials does not consider the thermal fluctuation of its subcomponents and thus only represents a material in a static equilibrium, which is unfortunately not the case of living matter. Then, a more realistic representation of a glassy material that exists near a glass transition is used; it consists in a soft matter approach with a mesoscopic description of thermal random fluctuations where the interactions among the material subcomponents are represented by a mean-field noise temperature parameter^66,67^. Soft glass matter theory may indeed describe adequately the rheological behavior of soft materials for which the classical elastic theory is deficient^68^, such as foams, emulsions, pastes, muds and, of course, living matter^69–72^. It indeed associates a probability to a modification of the macroscopic mechanical condition (obtained from a density of states and not an equation of states) and may be used to describe self-organization phenomena occurring in open systems that are out of equilibrium, like cells.

Cell culture experiments were carried out to determine the most appropriate model representing the mechanical behavior of the nucleus in the long term and to validate the H-VM proposed here. In particular, it was explored whether the model portrayed the nuclear strain dynamics correctly and revealed previous or stored information of its mechanical configuration, such as the magnitude of the nuclear deformation prior to the current state or even the stiffness of the microenvironment sensed by the nucleus in a previous state. This report demonstrates that the nucleus deformation of epithelial hepatic cells varies at least up to 5 days after being seeded. Also, it is shown that this dynamic behavior is very well described by the H-VM, which main assumption is that the nuclear strain caused by external mechanical cues, such as substrate stiffness is regulated by the mechanical properties of the structural components of the nucleus, as well as the dynamics of the network of filaments around it. From the statistical analysis of the experimental data, two different nuclear sizes are found in all conditions, also following distinct dynamic behaviors, which is more evident at higher stiffness and prolonged periods of time. Indeed, it was found for liver cells that the area projected by the nuclei is proportional to the substrate stiffness but their deformation is not. In time, the dynamic strain behavior of primary hepatocytes nuclei and a HepG2 cell line follows different behaviors that could be fitted with the H-VM. Furthermore, using the H-VM to fit experimental data, it is shown that it is possible to reveal a preexisting stable mechanical configuration of the nucleus, and even its long-term stiffness, caused by the mechanical properties of the substrate of the microenvironment perceived by the cell (a stable, reference configuration). Additionally, the dynamic of the nuclear Poisson’s ratio was measured in a plane transverse to the cell substrate. The nuclei of primary cells suffer a decrease in their Poisson’s ratio on both stiffness conditions over the time; however, for HepG2 cell line, a positive linear time-dependent Poisson’s ratio is found for the stiff condition while an auxetic behavior happens on the soft hydrogel after 36 hours in culture. These findings on the dynamics of nuclear strain and the variation of its Poisson’s ratio show that nuclear deformation is an irreversible process and suggest that the nuclear shape stores some information about previous mechanical states. With all these results, it is thus concluded that the H-VM model presented here enables a realistic simulation of the mechanical behavior of the cell nucleus at both short (seconds to minutes) and long (hours to days) timescales.

## Material and methods

### Fabrication of polyacrylamide hydrogels

Polyacrylamide hydrogels with specific elastic moduli were polymerized as described in^73^ and elastic properties were confirmed by microindentation assays using a FemtoTools FT-MTA-03, similarly to what was performed in^74^. Briefly, specific amounts of 40% acrylamide and 2% bis-acrylamide were mixed, added with 10% ammonium persulfate (APS) and 1% TEMED, and deposited onto 20 mm round glass coverslips which were treated previously with (3-Aminopropyl)triethoxysilane (APTES)/glutaraldehyde as mentioned in^75^; the polymerization reaction was carried out during 30 minutes at room temperature and the stiffness depends on the mixed quantities of the acrylamide and bis-acrylamide. The resulting polyacrylamide hydrogels were washed three times using 1x Dulbecco’s phosphate-buffered saline (DPBS); after washing, hydrogels were added with acrylic acid-NHS ester (0.02 g mL^−1^) and Irgacure 2959 (0.1g mL^−1^) mixed with rat tail collagen type 1 (0.1 g mL^−1^) dissolved in 20 mM acetic acid and exposed to UV light at 365 nm wavelength with a nominal power density of 3.3 mW cm^.2^ (UVP cross-linker CL-1000L) during 3.5 minutes. Hydrogels with the resulting conjugated protein were finally sterilized by washes with 1x DPBS plus antibiotics before seeding.

### Isolation of rat liver hepatocytes

The Animal Care and Use Committee of Facultad de Ciencias approved the animal protocol PI_2019_02_004 for isolation of rat liver hepatocytes on February 27th 2019. Fresh hepatocytes were isolated from livers of 250-300 grams Wistar rats, using a collagenase perfusion method described in^76^. After collagenase perfusion, isolated hepatocytes were separated by iso-density Percoll centrifugation in William’s E medium and cell viability was evaluated using trypan blue exclusion. Fresh viable hepatocytes were resuspended in attachment medium (DMEM-F12 supplemented with 4 mM GlutaMAX, 1 mM sodium pyruvate, 10 mM HEPES, 0.5 μg ml^−1^ amphotericin B, 1% insulin-transferrin-sodium selenite and 10% fetal bovine serum (FBS) and penicillin/streptomycin). Then, hepatocytes were cultured at a density of 2.5×10^5^ on round coverslips coated with polyacrylamide hydrogels of specific elastic modulus and conjugated with 0.1 mg/mL of rat tail collagen type I. Hepatocytes were allowed to adhere for 2 h in attachment media and replaced with feeding media (FBS-free attachment medium) afterwards. Simultaneously, hepatocytes were seeded on polystyrene plates covered with 1 mg mL^−1^ collagen dissolved in 20 mM acetic acid for standard control conditions.

#### Cell culture conditions

Hepatocytes were maintained in a feeding medium in standard culture conditions of 5% CO2 at 37°C for different durations as indicated, and media was replaced every 24 h. Human hepatoma HepG2 cell line was maintained in minimal essential medium (MEM) supplemented with 1 mM sodium pyruvate and 10 % FBS plus antibiotics. HepG2 cells were also cultured in standard culture conditions of 5% CO2 at 37°C.

### Immunofluorescence

Primary hepatocytes and HepG2 cells cultured on different stiffness conditions were washed with PBS 1X before fixation with 4% paraformaldehyde (PFA) at 37°C during 20 minutes. After fixation, cells were permeabilized using 0.1% Triton X-100 in PBS 1X by agitation at room temperature (RT) during 10 minutes and blocked with 10% horse serum in PBS 1X in agitation at RT for 1 hour; then samples were incubated with primary antibody against cytokeratin 18 (1:500) at 4°C overnight. Detection of primary antibody was performed by incubation with secondary antibody Alexa 594-coupled anti-mouse (1:500, Jackson ImmunoResearch, USA) during 1 hour at RT, and F-actin was stained using Alexa 488-coupled phalloidin (1:250, ThermoFisher Scientific, USA) for 1 hour at RT. Nuclei were detected by staining with 4’,6-diamidino-2-phenylindole (DAPI, 1:200, ThermoFisher Scientific, USA) for 10 minutes at RT, and after 3 washes with PBS 1X samples were mounted with Mowiol for preservation before imaging.

### Image acquisition

Samples were imaged by confocal laser scanning microscopy (Leica TCS SP8) and epifluorescence microscopy (Nikon Eclipse Ci-L). For confocal microscopy, hepatocytes were visualized using oil immersion 63X/1.40 NA objective while HepG2 cells were visualized using 40x/0.80 NA and 63x/0.90 NA water immersion objectives. Z stacking for nuclei (20 images of ~ 1 um) was acquired through blue channel, while CK-18 and F-actin were acquired through red and green channels, respectively. For epifluorescence microscopy, nuclei images were taken using 40X objective (5 fields per sample, blue channel). Image processing was carried out by using ImageJ software.

### Confocal microscopy image processing (segmentation and quantification of fluorescence intensity)

The images were analyzed in a Google Colab notebook using Python 3 language (programs are available upon request). Nuclei were segmented at different levels in the Z axis with the help of a moving average technique[^77^] and Otsu thresholding[^78^, ^79^], also a watershed algorithm[^80^, ^81^] allowed separation of objects that were in contact. To refine the object detection three filters were implemented: a number-of-objects filter, an area filter and a circularity[^82^] filter. Finally a contour classification was applied through a mean shift algorithm[^83,84^] in order to assign a set of objects (contours) to an unique nucleus. Information about area, perimeter, circularity and eccentricity[^80^] was computed for each located nucleus, additionally quantification of fluorescence intensity data was obtained for the red and green stacks using two concentric circles for each nucleus, the first circle is smaller than the biggest nucleus contour while the second circle is bigger than the biggest nucleus contour. This information was exported in “csv” and “png” files.

### Image processing of fluorescence images

The procedure was very similar to that implemented for the confocal image processing, but using only the blue channel. For the segmentation a weighted adaptive threshold[^85^] was implemented, followed by morphological operations for remotion of small objects and holes. Also two filters were used for segmentation refinement, one related to the area and one related to the circularity. Similarly to confocal processing, computation of the area, perimeter, circularity and eccentricity was done over the objects that remained in the last binary image and data was exported in “csv” and “png” files.

Check S3 for a detailed explanation of both image processing procedures.

### Data analysis

All the “csv” files generated by the segmentation software were analyzed with Wolfram Mathematica 12.1. Each file to analyze was sorted into an array of arrays. For the area, shape factor and eccentricity, the maximum value associated with each segmented nucleus was chosen. To calculate the height of the nucleus, the mean of each array was subtracted and the positions in the array of positive values were selected, since each position is related to a vertical Z value, a simple subtraction was performed. Interestingly, the values computed using this method agreed well with the value obtained manually using the ImageJ software. In order to analyze the data, different types of probability distributions were fitted for each specific condition. Due to the continuous nature of the phenomenon, only continuous distributions were used, such as: Normal, Log-Normal, among others. In the case of the mixed distribution, a deconvolution was performed to reveal the possible sub-populations in each condition. From these probability functions, the average and standard deviation were obtained and compared to the raw data values to verify whether the fit did not introduce artifacts or did not represent the mean of the raw data. Finally, from these data it was possible to calculate the deformation in a transverse and tangential plane to the nucleus, the aspect ratio, pre-strain and the Poisson’s ratio per condition.

### Statistical analysis

For the presented data, there are at least 2 independent experiments performed in duplicates. Statistical analyzes were performed with GraphPad Prism 8 and Wolfram Mathematica 12.1 software. All error bars reflect ± SD, comparisons among different conditions were performed by analysis of variance (one way ANOVA) with Tukey’s honestly significantly different post hoc tests with α = 0.05.

## Results

### Construction of the Hybrid-Viscoelastic Model (H-VM) for short and long timescales

#### Short timescale events are governed by viscoelastic behavior

The proposed mechanical model is based on a 1D viscoelastic description where the principal structural nuclear components were modeled as a linear viscoelastic material that approximated the viscoelastic properties reported for the NE (viscous lamin A/C *η*_*LA*_ and elastic lamin B *k*_*LB*_)^55,68,86^ and the Ch (*η*_*ch*_ and *k*_*ch*_)^28,87^ to a spring-dashpot system or Maxwell system (Figure 1A). These two Maxwell systems are connected in parallel with a long-range stiffness spring (*k*_*l*_) that represents the long-term stiffness, based on experimental evidence reported in the literature^88–90^. The possible relevance arises from the filaments network surrounding the nucleus, as recent reports have shown that cytoskeletal filaments (like intermediate filaments, keratins, vimentin and actin cap) play a relevant role in regulating the shape and mechanical properties of the cell nucleus^90–93^. The union of these elements forms a two-arm Maxwell generalized model (Figure 1B) and Equations 1 and 2 are thus the constitutive equations of the system:

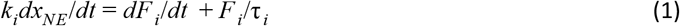

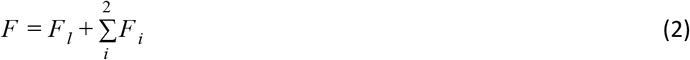

where *F* represents the total force exerted on the nucleus, *x*_*NE*_ is the deformation of the nuclear envelope, *F*_*i*_ represents the force on each arm, τ_*i*_ = η_*i*_/*k*_*i*_ is the relaxation time associated with each spring-dashpot subsystem and *F*_*l*_ = *k*_*l*_ *x*_*NE*_ is the force exerted on the long-term stiffness *k*_*l*_. To integrate the contribution of the nuclear pore complexes to the total nuclear deformation, the φ = *x*_*p*_/*x*_*NE*_ parameter was defined as the ratio between the deformation *x*_*p*_ of pores and the deformation *x*_*NE*_ of the nuclear envelope, hence obtaining Equation 3 that describes the total deformation of the nucleus *x*_*n*_.

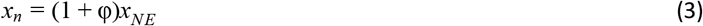

**Figure 1.**
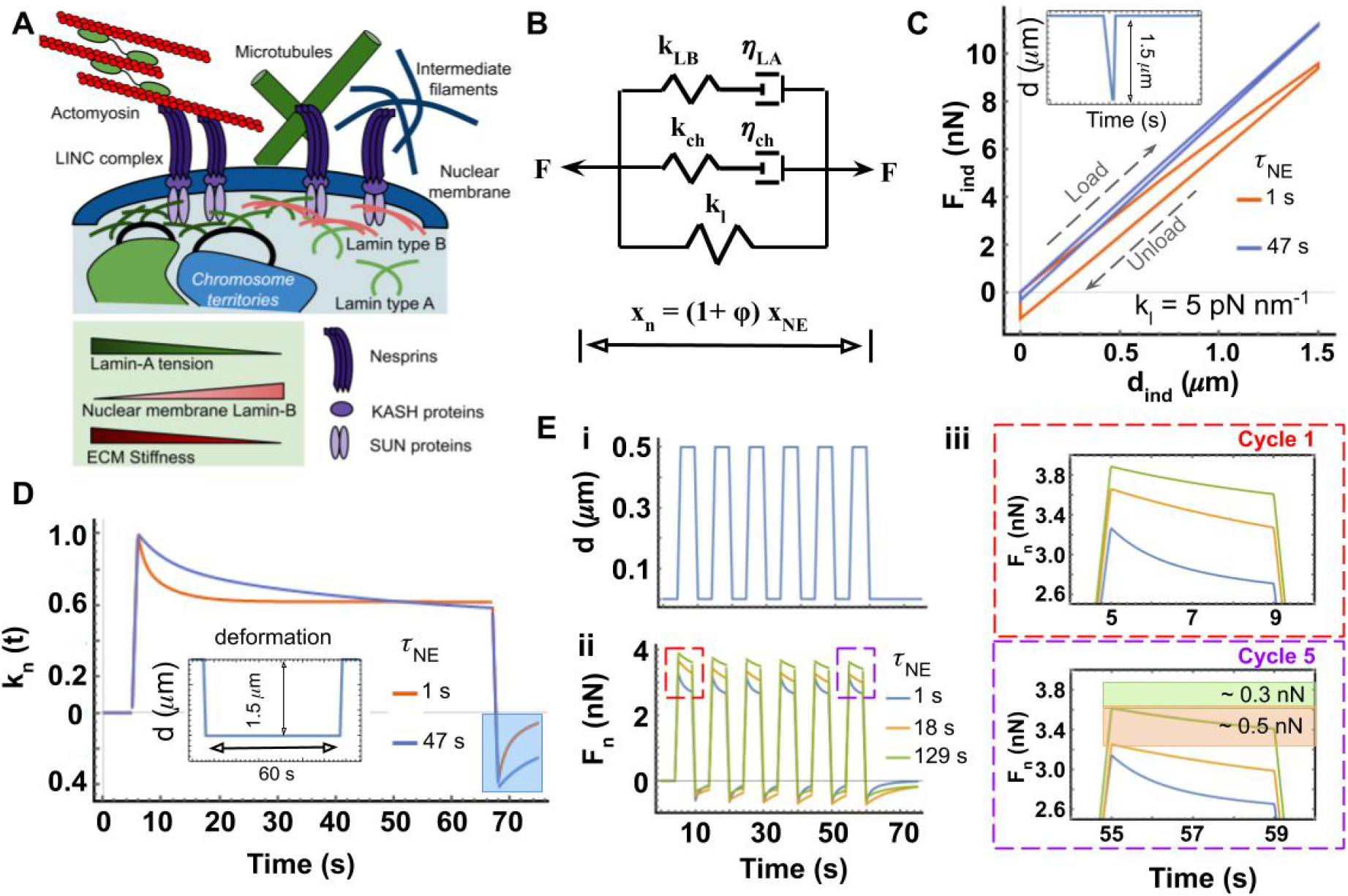
Viscoelasticity governs short timescale mechanical nuclear behavior. A) The principal subcellular components that determine the mechanical properties of the nucleus are: the nuclear envelope (NE), the acetylation state of chromatin (Ch) and the dynamic reorganization of cytoskeleton governed by the intermediate filaments, microtubules and the actomyosin apparatus, which behavior is highly influenced by the stiffness of the extracellular matrix (ECM). B) The hybrid viscoelastic model (H-VM) is proposed based on a two-arm generalized Maxwell model recapitulating the rheological behavior of the nucleus together with the NE activity and the dynamics reorganization of the cell cytoskeleton, assuming η_*LA*_ = η_*LA*_(*t*), *k*_*LB*_ = *k*_*LB*_(*t*) and *k*_*l*_ = *k*_*l*_(*t*). However, at short (seconds and minutes) timescales they are considered constant. C) Simulation of a force-deformation curve for an indentation assay on a nucleus modeled by H-VM. An indentation depth of 1.5 μm and a 0.5 s delay are used with a load velocity of 1 μm/s and an unload velocity of 3 μm/s. Two NE relaxation time (defined as τ_*NE*_ = η_*LA*_/*k*_*LB*_) were modeled. D) Simulation of a relaxation assay for a modeled nucleus (indentation at 5 s at a 1.5 μm depth and a velocity of 1.5μm/s). The deformation is maintained constant for 60 s and liberated afterwards (same velocity). E) Simulation of (i) a periodic deformation of the nucleus with a 10 s period with 4 s pulses and a deformation of 0.5 μm and a load/unload velocity of 0.5 μm/s; (ii) response to the deformation, force in nN; (iii) zoom of cycles 1 and 5 where a decrease of 0.3 - 0.5 nN is observed, associated to nuclear stiffening.

The principal assumption of the H-VM is that the parameters η_*LA*_, *k*_*LB*_ and *k*_*L*_, considered constant in the generalized Maxwell model, are now dependent on the tension *T* and time *t*, hence expressed as: η_*LA*_ = η_*LA*_(*T*, *t*), *k*_*LB*_ = *k*_*LB*_(*T*, *t*) and *k*_*l*_ = *k*_*l*_(*T*, *t*). These equations, described further in section 2.1.2, hence illustrate the active mechanical behavior of the nuclear envelope and either the glassy dynamics or hyperelastic behavior of the filaments network surrounding the cell nucleus. Furthermore, the nuclear deformation is calculated by assuming either a linear or nonlinear dynamic behavior of *k*_*l*_(*T*, *t*) and a stable, reference configuration caused by the contractility *f*_0_. The parameter *k*_*l*0_ ≡ *k*_*l*_(*t*,*f*_0_) represents this basal mechanical state of the nucleus caused by external mechanical stimuli and, from this stable state, the variation of the mechanical properties of the nucleus is defined at large time scales. The parameters η_*ch*_ and *k*_*ch*_ are considered constant in the long timescale because it has been reported that the chromatin only supports small deformation (below 30 %), unlike the NE that supports large deformations^28,94,95^.

Using the software Wolfram Mathematica 12.1, a dynamic solver of Equation 1 and 2 was programmed to simulate the mechanical response for three common mechanical tests: microindentation, cyclic/periodic dynamic deformation assays and relaxation tests. Because these mechanical analyses are commonly performed in less than a few minutes, the parameters η_*LA*_, *k*_*LB*_ and *k*_*l*_ are constant in the time of the experiments; indeed, they vary only in long timescales (typically hours, as will be shown further) and they are impacted by the ECM stiffness^55,96–98^. An interactive application is available on demand (NOTE for reviewers: it will be posted on a GitHub repository upon acceptance of the paper and is available to you for review), with a complete description of each parameter to allow the reader to explore a broad range of values and combinations of those. Panels C, D and E in Figure 1 present a few mechanical observations taken from the literature that were satisfactorily approximated by our model.

In Figure 1C, the simulation of a classic indentation assay is shown. The two NE relaxation times selected in our simulations are associated with that corresponding to an hepatocyte on a soft tissue τ _*NE*_ = 1 *s* and with the cutoff value reported for liver fibrosis τ _*NE*_ = 47 *s* ^55^ (Table S1). It may be observed that by increasing the NE relaxation time τ _*NE*_ = η_*LA*_/*k*_*LB*_, there is an increase of the slope in the force-displacement curve. This is commonly interpreted as an increase in the elastic modulus of the material under test. Moreover, the graph presented in Figure 1C displays a linear behavior of force vs. distance with a shift between load and unload slopes, different from the one expected for an elastic material, following the *d*^3/2^ behavior of the Hertz model^99^. Interestingly, this simulated force vs. distance curve has already been described in the literature: Schäpe *et. al.* showed that when lamin A expression is increased in the NE of the nucleus of *Xenopus oocytes*, the slope of the indentation curve obtained with an atomic force microscope (AFM) also increases and presents a linear behavior.^31^. This response was interpreted back then, in a classic elastic approach, by a greater Young’s modulus. However, in our model, Equations 1 and 2 show that it is most probably caused by an increase of the viscosity associated with a gain in lamin A, assuming that this increment only modifies the viscosity of NE without perturbing the other components. The Ch relaxation time and the long-range stiffness were kept constant and typical values of τ _*ch*_ = 5 *s* and *k*_*l*_ = 5 *pNnm*^−1^ were chosen (see Table S1). All this thus suggests that the apparent increase of the curve slope measured with AFM (first accounted on the elastic modulus) may be due to the fact that the time taken to perform the mechanical characterization using indentation is similar to the relaxation times of NE and Ch. Again, in the interactive *Mathematica* application available on demand (NOTE for reviewers: it will be posted on a GitHub repository upon acceptance of the paper and is available to you for review), it is possible to explore the impact of indentation velocity and variations of the other parameters, such as: slope of the force vs. displacement curve). τ _*NE*_, *k*_*l*_ and τ _*ch*_ on the value of the nuclear stiffness (*i.e*, the slope of the force vs. displacement curve).

The relaxation modulus *k*(t) (commonly known as *E*(t) when the test is performed on a 3D material) is defined as the stress variations under a constant strain. It represents the variation of the force per unit area under a constant deformation and the purpose of the relaxation test is to reveal the elastic and viscous properties of the material in the timescale in which it was performed^100,101^. It is important that a model that seeks to describe the nuclear mechanics accurately is able to fit or predict the relaxation module of the cell nucleus *k*_*n*_(*t*). In order to observe the impact of the NE viscoelastic behavior on *k*_*n*_(*t*), only the values associated with NE were modified, since it has been reported that the levels of lamin A/C and B1 depend on the ECM stiffness^32,55^. The simulation results are presented in Figure 1D for a constant compression of the nucleus of 1.5 μm during 60 seconds. These parameters are standard for a relaxation assay; 1.5 μm complies with the Hertz model hypothesis of small deformations (of less than 30% of the nucleus radius assuming a 10 μm height for hepatocytes)^102^ and 60 seconds is common to determine the magnitude of relaxation times in such assays. The time response that was obtained is comparable to that presented by Yin and collaborators^100^, with an almost instantaneous load and a recoil (shadowed in a blue box) after liberation of approximately 40%. In this assay, the direct influence of the NE relaxation time is also clearly visible on the relaxation of the complete nucleus. Finally, Figure 1E shows the mechanical response (Figure 1E-ii) of the nucleus to a cyclic deformation of 0.5μm with a 10s period (Figure 1E-i). The NE relaxation times used in this assay are τ _*NE*_ = 1s, τ _*NE*_ = 18s and τ _*NE*_ = 129s and correspond respectively to healthy liver, healthy lung and highly fibrotic liver tissues (see Supporting Information S1). Plots presented in Figure 1E-iii are zooms of the maximal values of cycle 1 and 5 of Figure 1E-ii. By comparing the force in these two cycles, separated by more than 40s in time, it is observed that for τ _*NE*_ values greater than the duration of the deformation pulse, the force slowly decreases as the number of cycles increases (a reduction of approximately 0.5 nN is observed for τ _*NE*_ = 18 *s* and 0.3 nN for a greater relaxation time of 129 s) while the magnitude of the force remains constant over time if τ _*NE*_ is lower than the duration of a deformation pulse. Since the magnitude of the cyclic deformation remains constant, it may then be inferred that this reduction of the magnitude of the force illustrates a possible nuclear stiffening, dependent on the viscoelastic characteristics of NE and Ch state. This interesting phenomenon has already been described in isolated HeLa cells by Guilluy *et. al.^11^* but found absent in nuclei of lamin A/C knockdown cells, even after several cycles, thus suggesting that this phenomenon is principally determined by the mechanics of lamin A/C, or τ _*NE*_, as shown now in our model.

Importantly, the results of the H-VM shown in panels 1C, 1D y 1E suggest that the viscosity of the NE, which levels are generally associated with the tissue stiffness^55^, is the main determinant of the nuclear mechanical response at short time-scale.

After an exploration of the impact of the *k*_*l*_ value on the exerted force using the aforementioned application, it is observed that the value of this stiffness only modifies the magnitude of the response without altering the temporal behavior at short timescales (in the range of 0-300 s). Additionally, the *η*_*ch*_ and *k*_*ch*_ parameters were considered constant, since the dynamic behavior of the viscoelastic response is probably due to the condensation (or decondensation) of chromatin^103^. These simulations with constant parameters reveal that when mechanical tests are performed with a measurement time of the same order of magnitude as the relaxation time of the NE or Ch (~ 0-300 s) the results obtained in the experiments are likely to be affected by the viscous contribution of the nuclear structures even though they are usually not acknowledged. Therefore, for a correct interpretation of this type of measurement, the viscoelastic properties of the nucleus need to be recognized. However, for the phenomena occurring at longer timescales, similar to those of the replacement of the nuclear lamins of the NE and histones of the Ch (typically in the order of dozens of hours^104^), it is necessary to include the active response of the NE and the influence of the reorganization of the cytoskeleton on the mechanical response of the nucleus. This is addressed in the following section.

### Long timescale events are governed by NE activity and glassy behavior

Although most mechanotransduction processes occur in a short period of time (seconds and minutes)^13,105^, some irreversible phenomena such as aging^106^ and mechanical memory^107^ (mechanisms allowing the cell to store information on a particular mechanical condition for a long period of time), occur at long timescales from days to years, in the context of the cell nucleus. The cell nucleus is commonly modeled as an elastic or hyperelastic material and it successfully describes cell migration^108^ and the impact of spatial confinement on nuclear architecture^65^ but the models are limited when aiming at long-term events like plasticity, mechanical irreversibility and aging, because the specifications of the model do not overlook dissipative effects^11,33,46,109–112^. These particular events typically lie in timescales beyond the characteristic time of the viscous dissipation of the nuclear envelope and chromatin acetylation state^54,55,108^. For these long timescales, it may be hypothesized that the observed irreversibility originates from the active behavior of the nuclear envelope and from the continuous reorganization of the cytoskeleton. Mechanically, these two phenomena are indeed known to confer structural stability to the nucleus through the dissipation of the forces that deform it. First, the activity of the nuclear envelope consists of the regulation of its mechanical properties to maintain the structural integrity of the nucleus^113,114^. Additionally, the reorganization of the cytoskeletal filament network that surrounds the nucleus depends not only on static mechanical stimuli originated by the tissue stiffness ^88,92,98^, but also on dynamic mechanical stimuli such as periodic deformations caused by external forces^109,115^.

In order to model these phenomena, it was assumed that the parameters η_*LA*_, *k*_*LB*_ and *k*_*l*_ in the Equations 1 and 2 all become tension and time dependent, so we assume the following relations: η_*LA*_ = η_*LA*_(*T*, *t*), *k*_*LB*_ = *k*_*LB*_ (*T*, *t*) and *k*_*l*_ = *k*_*l*_ (*T*, *t*). In view of the dependence of the levels of expression of nuclear lamins with the extracellular matrix (ECM) stiffness^55^ and of the inhibition of the phosphorylation of those lamins (especially lamin A/C) caused by the mechanical tension of the nuclear envelope^116^, a gene circuit has been incorporated in our model (Figure 2A), based on one that was proposed elsewhere^117^. Its main purpose is to account for the dynamic regulation of the synthesis, accumulation and degradation of lamin A and B1 as a response to the tissue stiffness. The process by which the explicit form of these equations was found is presented below.

**Figure 2.**
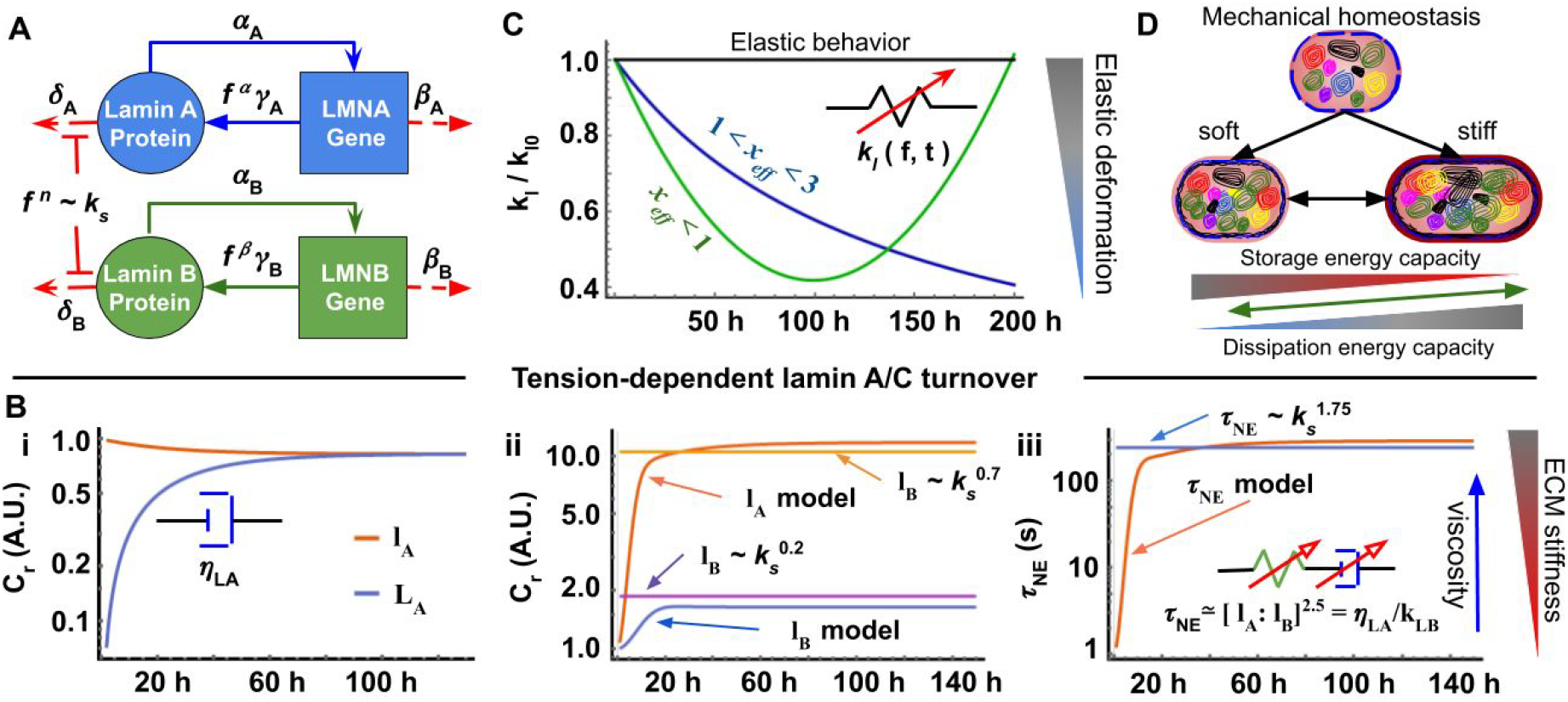
Nuclear envelope (NE) activity and cytoskeletal organization govern long time mechanical nuclear behavior. A) Representation of the genes circuit defined by a system of differential equations depicting the activity of lamins A/C and B1 that form the NE with the coefficients *α*, ⍰, δ and γ being the synthesis and degradation coefficients and *L*_*i*_ and *l*_*i*_ the concentrations of mRNA and proteins respectively. These coefficients are associated with the corresponding half-lives *t*_*1/2*_ of the protein and mRNA to simulate. The equations describe the inhibition of protein degradation and the increase of mRNA synthesis due to the tension suffered by the protein and caused by a change in the mechanical state of the system. B) (i) Rate of replacement of lamin A in a stable mechanical state, calculated in the particular case of a primary hepatocyte. (ii) Increase of protein synthesis and inhibition of protein degradation due to a tension increase caused by a greater ECM stiffness for an hepatocyte. (iii) Mechanical response associated to the increase of the ECM causing an increase of τ _*NE*_. C) Temporal evolution of *k*_*l*_ when modeled as an elastic material (horizontal black curve) or a glassy material close to its glass transition temperature *x*_*g*_ ≡ 1. In the latter case, the glassy behavior is defined by the mean field temperature parameter *x*_*eff*_. The blue curve represents a weak power-law when *x*_*eff*_ ∋ (1, 3) and the green curve represents a pseudo power law when *x*_*eff*_ < 1 with a stable point close to 100 hours. The value *k*_*l*_ (*t* = 0) = *k*_*l*0_ is associated with the reference mechanical state of the system. D) Summary of the long-time impact of τ _*NE*_ and *k*_*l*_ assuming that *k*_*l*_ follows a weak power-law behavior. When the nucleus mechanical state is perturbed, its response depends on whether the current microenvironment is softer or stiffer than the former one. If softer, it tends to acquire a more elastic and less deformable behavior, while a stiffer substrate creates tension and leads to a more dissipative and less deformable nucleus.

The system of four equations which describes the gene circuit is presented below:

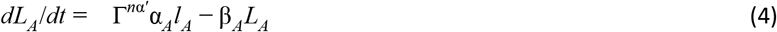

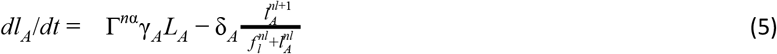

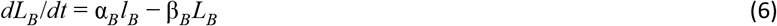

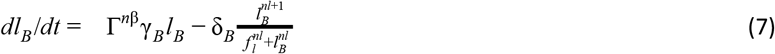

where *L*_*A*_, *L*_*B*_ y *l*_*A*_, *l*_*B*_ represent the concentrations of mRNAs and proteins of lamina A and lamina B1 respectively, with their synthesis (α_*i*_ y γ_*i*_) and degradation (β_*i*_ *y* δ_*i*_) rates dictating their expression levels and modifying their mechanical influence on nuclear deformation. The parameter *f*_*l*_ = *f*_*r*_ − *f*_0_ / *f*_0_ represents the relative change of the tension generated by the actomyosin apparatus contractility and is thus associated with the ECM stiffness. Therefore, *f*_*r*_ is the effective tension exerted on the nucleus with respect to an equilibrium state *f*_0_, which is defined as the intrinsic nucleus tension in a stable mechanical state, which is in turn associated with a reference nuclear size due to the cytoskeletal stress of a given previous state. The coefficient *nl* is defined as the cooperative coefficient (≥2) which is typical of multimeric interactions^59^. Additionally, the parameter Γ^*ni*^ = *f*_*r*_ / *f*_0_ is the tension ratio and describes the increment of the rate of synthesis as a function of the relative tension; the coefficient *ni* thus represents the tension-dependent synthesis level.

The gene circuit is set up with the following half lives values for nuclear lamins and their associated mRNAs, according to the literature^104^: *t*_1/2_ = 89.46 ± 14.85 *h* for lamin A and *t*_1/2_ = 114.16 ± 18.95 *h* for lamin B1, *t*_1/2_ = 18.79 ± 3.12 *h* for mRNAs associated to lamin A and *t*_1/2_ = 7.14. ± 1.19 *h* for mRNAs associated with lamin B1. Considering that a cell located in its natural microenvironment does not modify its mechanical properties spontaneously, a mechanical homeostasis state may be defined. This stable mechanical state fulfills the following conditions: it is almost constant over timescales larger than the turnover of the proteins constituting the structural components of the cell, it is specific to each cell type and it is stable with the mechanical cues of its corresponding environment. For any given cell, it may be hypothesized that this reference state coincides with the last stable mechanical state of the cell. Interestingly, it is shown below that the stiffness of the original soft tissue microenvironment is traced back for liver primary cells, and a much stiffer reference state is found for a hepatocellular carcinoma cell line. Therefore, *f* _0_ and the initial conditions of the protein levels of the Equation 5 and 7 establish a reference state for the adherent cells, for which the nuclear proteins levels are tension dependent. The Equations 4, 5, 6 and 7 were solved numerically using Wolfram mathematica 12.1 and the solver is available for the reader as an online application (link in the Supporting Information supplementary material 3) with a description of the base parameters. Figure 2B-i shows the lamin A levels (both protein *l*_*A*_ and mRNA *L*_*A*_ levels). They were calculated with the proposed gene circuit and for the particular case of a primary hepatocyte coming from a natural stiffness of 0.75 - 1 kPa (representing a cell from a healthy liver, in the homeostatic state, as reported in the literature^118^). It can be observed that these levels reach a stable state, *i.e.* end up being constant over time. In contrast, Figure 2B-ii shows a gradual increase in the accumulation of both *l*_*A*_ and *l*_*B*_ protein levels, until they finally reach a different stable state. This increase of synthesis and inhibition of protein degradation is thought to represent an increase in tension, a response of the hepatocytes when suffering an increase in their substrate stiffness, simulating a culture on a 23 kPa material. Hence, to simulate this increment in time of the contractility generated by the actomyosin apparatus (until it reaches another stable tension level in response to the increment of the substrate stiffness), it is assumed that *f* behaves as an accumulation function of a logistic distribution. This is justified by the fact that it was reported that this kind of S-shape function may capture adequately the stress-strain response of a macroscale elastic network in a viscous medium (like the cytoskeleton) at short and long time ranges (thresholds)^119^. So, the plot presented in the Figure 2B-ii is possible to observe the rapid increment of the protein concentration until reaching a constant value which means that the tensión exerted on the NE reaches an equilibrium state. Additionally, in order to be consistent with the experimental data reported by Swift. J. *et. al.* in^55^ 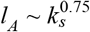 and 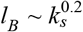. Figure 2B-iii shows that the accumulation of lamin A and lamin B1 will be experimentally reflected by an increase of the nuclear envelope relaxation time τ _*NE*_. To obtain this, in order to associate the protein levels with the relaxation time, Swift *et. al.* have proposed the following relation^55^: τ _*NE*_ ≃ [*l*_*A*_: *l*_*B*_]^2.5^ ≃ η_*LA*_/*k*_*LB*_ from which the following is obtained:

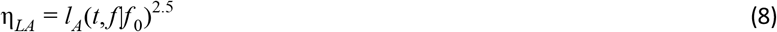

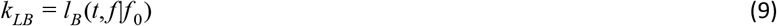

were *l*_*A*_(*t*, *f* |*f*_0_) and *l*_*B*_(*t*, *f* |*f*_0_) are the solutions to the Equations 4 - 7 and are the explicit form of the equations η_*LA*_ = η_*LA*_(*T*, *t*), *k*_*LB*_ = *k*_*LB*_(*T*, *t*). It is thus possible to infer an important direct relationship between the internal NE viscoelasticity (relaxation) and the external ECM stiffness *k*_*s*_, since the solution of the Equation 4, 5, 6 and 7 provides the relations 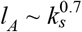 and 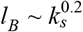 as seen in Figure 2B-ii and in the interactive *Mathematica* application available on demand (NOTE for reviewers: it will be posted on a GitHub repository upon acceptance of the paper and is available to you for review),.

In order to elucidate the explicit form of *k*_*l*_ as a tension- and time-dependent parameter *k*_*l*_ = *k*_*l*_(*T*, *t*), it was hypothesized that this long-term accounts for the behavior of the network of filaments that surround the nucleus. Due to its rheological behavior, the soft glass matter theory described by Sollich P. et al.^67^ was used here. The basic assumption is that the thermal movement on its own is not sufficient to describe a complete structural relaxation and the resulting mechanical behavior is thus a natural consequence of two of the intrinsic suppositions of the soft glass framework: structural disorder and metastability of the subcomponents^66–68,109^. A finite mesoscopic system where each subcomponent of the system is a filament that is part of the network surrounding the cell nucleus was thus assumed here. Each filament presents a linear elastic behavior described by a constant *k*_*i*_ and a deformation *l*_*i*_ with respect to the equilibrium length *k*_*i*0_. Hence, the elastic energy *E*_*i*_ associated to the i^th^ filament in a metastable state may be given by 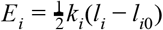. To model the effects of structural disorder, the total elastic energy of the system is described by a density of energy states *E*_*T*_. Then, the state of a macroscopic sample is characterized by a probability distribution *P* (*E*_*T*_, *l*; *t*), where *l* is the average strain of the network. Besides, in the context of the mean-field theories, the interactions among the filaments are subsumed into an effective noise temperature parameter *x*_*eff*_. The form of this probability distribution in a equilibrium state *P*_*eq*_ thus depends on the parameter *x*_*eff*_ and since the probability density function may take the form *P*_*eq*_(*E*) ~ *exp*(*E*_*T*_ /*x*_*eff*_) ρ(*E*_*T*_) all this leads to a power law for 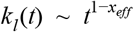 in the long time range, or low frequencies (see ^67,120^ for a more detailed explanation). Thus, only the cases when 1 < *x*_*eff*_ < 3 and *x*_*eff*_ < 1 will be taken into account. Indeed, we have:

- when *x*_*eff*_ > 3, *k*_*l*_ represents a Maxwell system (that represents a fluid-like behavior for which the nucleus acquires a purely viscous behavior that is not consistent with experimental observations).
- In the (3,2) interval, the behavior of *k*_*l*_ is described by a power law and in the (2,1) interval by a pseudo power law or a weak power law.
- Finally, when *x*_*eff*_ < 1 then *k*_*l*_ tends to a non-stationary state^67^.

Additionally, *k*_*l*_(*t*) must comply with the fact that, at short periods of time it remains almost constant, and then: 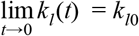. Then, for the case when 1 < *x*_*eff*_ < 3, if we assume that the *k*_*i*_ constants associated with the i^th^ filament are different and acquire random positive values, without loss of generality it can be assumed that the energy density function follows a gamma distribution, and therefore:

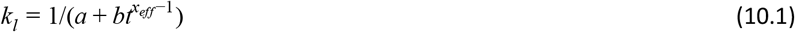

where *a*, *b* and *x*_*eff*_ are constant parameters that may be found by fitting experimental data, since equation 10.1 satisfies 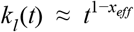 and 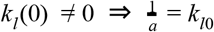.

On the other hand, when *x*_*eff*_ < 1, it is implied that *x*_*eff*_ < *x*_*g*_ (where *x*_*g*_ is the glass transition temperature and is set up to 1). Consequently, a state of equilibrium does not exist since the probability to access all possible energy states decreases and the system shows various aging phenomena; *i.e. P* (*E*_*T*_, *l*; *t*) does not present a steady state in time. However, if a high energy cutoff is imposed on the energy density function, 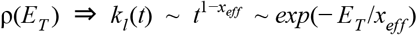, leading to the following equation using a third order Taylor expansion at *x*_*g*_ = 1 :

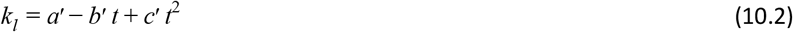

where the constants *a’*, *b’* and *c’* can be found by fitting experimental data and we also obtain that *k*_*l*_(*t* = 0) = *a* = *k*_*l*0_.

In Figure 2C, a comparison between a glassy dynamic regime and a hyperelastic behavior that is constant over time, is presented. The graph shows the temporal evolution of *k*_*l*_ when modeled as an elastic material (horizontal black curve) vs. a glassy material close to its glass transition temperature. In the latter, the glassy behavior is defined by the mean field temperature parameter *x*_*eff*_. The blue curve represents a weak power-law when 1 < *x*_*eff*_ < 3 and the green curve represents a nonlinear function when *x*_*eff*_ < 1 with a stable point close to 100 hours. The value *k*_*l*_ (*t* = 0) = *k*_*l*0_ is associated with the reference mechanical state of the system and was set to 1 in all cases. the coefficients *b, b’* and *c’* of Equations 10.1 and 10.2 may be associated with the half-life of the cytoskeletal proteins in a range of 50-120 hours (known typical half-lives of keratins, vimentin and other cytoskeletal proteins filaments^97,121^). As expected, the blue curve tends to reach a stable state that depends on the value of coefficient *b*, while the curve that represents Equation 10.2 presents a minimum that depends on the values of coefficients *b’* and *c’*. For a deeper exploration and understanding of these equations, it is possible for the reader to inspect the influence of the parameters *a, b, a’, b’, c’* and *x*_*eff*_ on the behavior of Equations 10.1 and 10.2, using the application developed for this work and mentioned above.

Finally, it is considered that Equations 1–3 and 8-10.1 (or 10.2) fully describe the Hybrid Viscoelastic Model (H-VM) of the cell nucleus and Figure 2D summarizes the stable nuclear behavior at greater timescales of hours and days. Nuclei of cells that are cultured on substrates with a stiffness comparable to that of their natural/original mechanical homeostasis state do present an almost constant level of expression of lamins A and B1 and *k*_*l*_ remains invariant as well. However, if the nucleus is perturbed by a tension greater than the one corresponding to its mechanical homeostasis, the levels of expression of nuclear lamins increase (Equation 8 and 9) and the value of *k*_*l*_ may either monotonously decrease until it reaches a minimum stable value (when Equation 10.1 is valid), or once it reaches a minimum increase its elastic energy storage capacity again (in case of Equation 10.2). This is of particular importance because, essentially, it implies an increased dissipative capability and deformability of the nucleus on stiffer substrates or when subjected to larger deformations, while softer materials lead to an increase in elastic deformation capacity (low strain regime) and a lower dissipative capability.

A direct consequence of H-VM is that if the nucleus deformation at long timescales is described by Equation 10.2 instead of Equation 10.1, the nucleus recovers certain elastic deformation capacity through reorganization of the filament network around it. This reorganization process has an energy cost for the cell, since it is not a completely reversible process. Then it is suggested that the cell nucleus has gone through a long-term deformation process that may be a form of aging, the origin of which lies in the irreversibility nature of nuclear deformation. To define the correct Equations 10 to describe the behavior, experimental data are required and the best fit is selected (see the experimental section).

### Cell nucleus strain simulation

Simulations presented in Figure 1 and obtained from the H-VM model in one dimension were suited to reproduce theoretically the nuclear behavior reported in the literature with indentation assays^31^, relaxation^100^ and periodic deformations^11^ tests. Additionally, an explicit expression was proposed (Equation 8 and 9) to model the active behavior of the NE^55,116^ and the glassy dynamics of the filaments surrounding the nuclear structure^68^. Although it is a simplified 1D model, it proved consistent with a 3D FEA simulation (See Supporting information S2). In order to validate whether the H-VM is adequately connectable to the other mechanical elements of a cell and help describe better the mechanical response of the nucleus to physiological forces exerted at long timescales, the mechanism of cell spreading induced by substrate stiffness was tested. Given the nature of this phenomenon, it may be well approximated by a 1D model (**Figure 3A**), because it is caused by a direct link between the ECM and the nucleus: during cell spreading, the nuclear deformation is mainly due to a contractile tension produced by the actomyosin apparatus and dependent of the substrate stiffness^116^. In order to simulate this phenomenon, the representation of the mechanical chain of transmission of Figure 3B is proposed. This system depicts the union between the H-VM model and a set of mechanical elements that was proposed earlier by Shenoy’s group^64,65^ and simulates the chemo-mechanical activity ϱ of the myosin molecular motors in parallel with the resistance to compression *k*_*μT*_ of the microtubules, the tension of actin filaments *k*_*a*_ and the effective stiffness *k*_*eff*_ of the adhesion complex united to the ECM with a stiffness (please see Supporting Information for a detailed description and explanation of each element). Panel 3C presents the dissipation of the actomyosin stress as a function of the levels of nuclear lamin at different times, corresponding to the relaxation time of the Ch (5s) and the NE (129s) taken for an hepatocyte inside a fibrotic liver of 23 kPa^122^ (See Table S1). It clearly shows that the magnitude of the stress σ(*t*) generated by the actomyosin apparatus depends on the levels of η_*LA*_ and *k*_*LB*_ and therefore on time. For higher values of η_*LA*_ and *k*_*LB*_, the initial magnitude of the stress is maintained for a longer period of time but undergoes greater dissipation once it reaches a stable state after a time τ _*NE*_ = η_*LA*_/*k*_*LB*_, which depends on the substrate stiffness *ks* and long-term nucleus stiffness *k*_*l*_ (t) (see Figure S1A). The white (τ _*NE*_ = 1 *s*) and red (τ _*NE*_ = 129 *s*) dots represent the relaxation times of a healthy and a highly fibrotic liver, respectively (see table S1). The nuclear deformation *ε*_n_(t) was calculated using the Boltzmann integral (Equation 11, below) that is a function of the creep modulus *J(t)*, which temporal behavior is presented in panel 3D and depends mainly on the values of η_*LA*_(*t,f*), *k*_*LB*_(*t,f*) and *k*_*l*_(t), then of the ECM stiffness *k*_*s*_. As can be seen in this panel, *J(t)* is inversely proportional to the value of the ECM stiffness, which is consistent with previous reports in the literature^55,103^. Equation 11 describes the time-dependent strain that results from the application of a time dependent stress σ(*t*) with a stable value that is a function of the substrate stiffness (Figure S1A); and in the equation, the parameter ξ is a convolution variable:

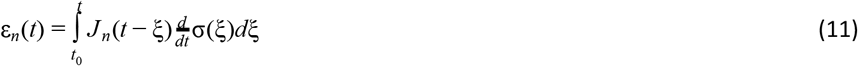

**Figure 3.**
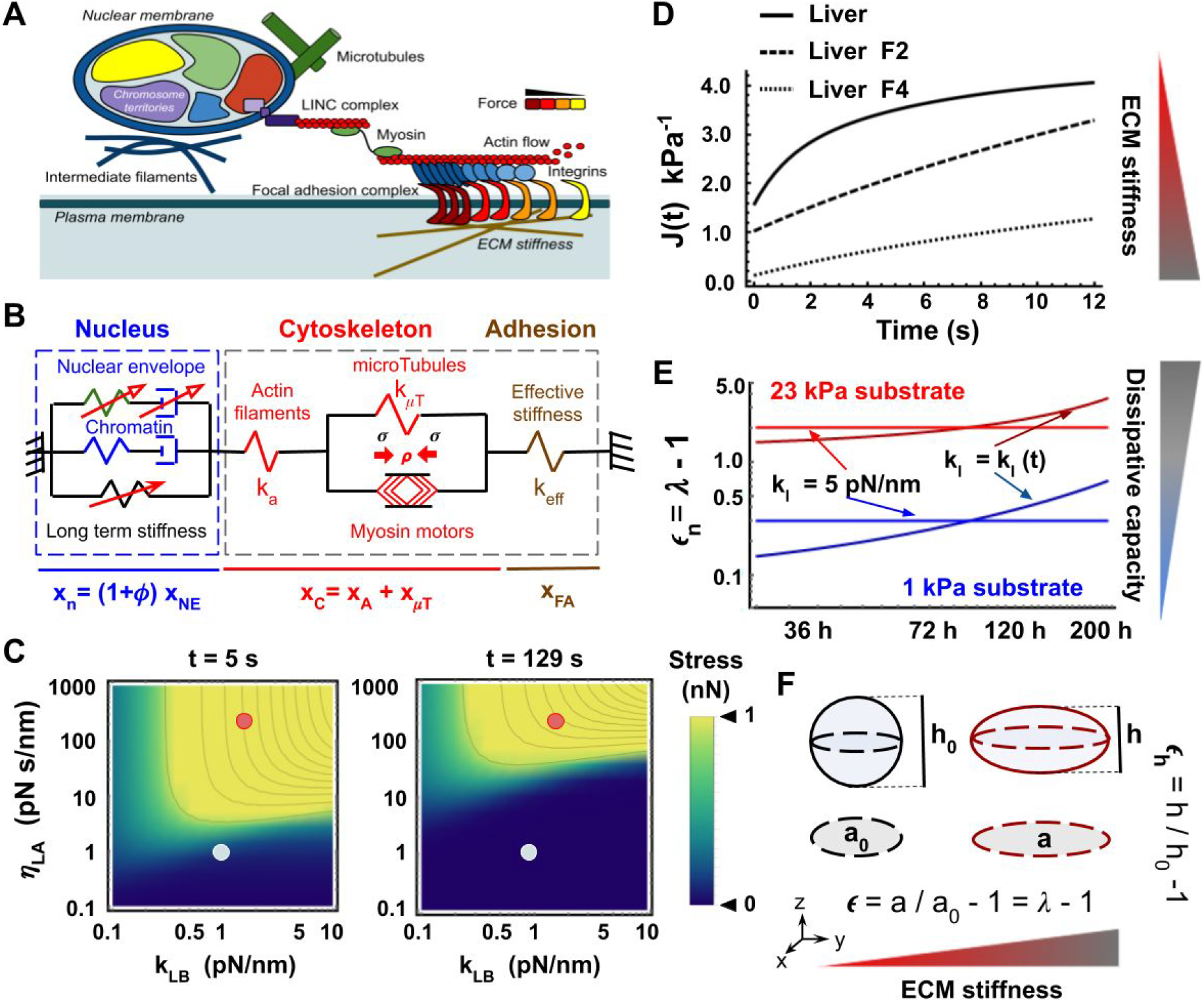
Mechanical representation of an adherent cell. A) Schematic diagram of the direct link between ECM and nucleus via the adhesion complex and actomyosin apparatus, taking the resistance to compression into account of the microtubules and the intermediate filaments network surrounding the nucleus, providing structural support. B) Mechanical representation of an adherent cell using the H-VM model. The red arrows in the nucleus box, on the representative elements of the nuclear envelope and the long-term stiffness depict the temporal dependency of these parameters. It is also considered that focal adhesions are in dynamic equilibrium (independent of time). C) Dissipation of the actomyosin stress as a function of nuclear lamin levels at t=5 s (relaxation time of the chromatin Ch) and at t=129 s (relaxation time of the NE for an hepatocyte inside a highly fibrotic liver of 23 kPa^122^ see Table S1). The white dots represent the lamin A/C and lamin B1 levels of a healthy hepatocyte (soft liver tissue) and the red dots show the same levels for an hepatocyte originating from a liver in a state of stage 4 fibrosis (F4). D) Fluence modulus of the H-VM model as a function of the ECM stiffness for a soft, healthy liver (~1 kPa), a stage 2 fibrotic (F2) liver (~12.5 kPa) and a a stage 4 fibrotic (F4) liver (~ 23 kPa). The magnitude of the fluence decreases with an increasing ECM stiffness. E) Nuclear strain as a function of time for two different models for the nucleus: an elastic nucleus (horizontal line) and a glassy solid nucleus vítreo (ascending line) are shown. The larger the ECM stiffness, the greater the deformation and capacity to dissipate the elastic energy (Equation 10.1 shows that kl(t) decreases with time. F) Projected area used to calculate the longitudinal nuclear deformation *ϵ* and the deformation of the nuclear height. The reference values of projected area a0 and height h0 come from a stable mechanical condition.

In Equations 1–3, no particular explicit reference system was considered. However, one had to be imposed by the equations that govern the temporal behavior at longer timescales, *i.e.* Equations 8, 9 and 10.1 (or 10.2). Indeed, Equation 11 is defined for times greater than *t*_0_ ≥ 0 where *t*_*0*_ represents a reference stable state. Therefore, the total nuclear strain given a reference condition is calculated using Equation 12:

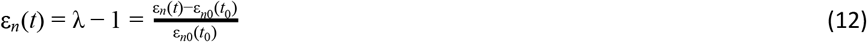

where λ = *l*/*l*_0_ is the common definition of strain, ε_*n*_(*t*) is the Cauchy strain definition and ε_*n*0_(*t*_0_) = *l*_0_ is the configuration of a reference stable state. This full system of equations (see Supporting Information S1) was solved in the Laplace domain, for simplicity, using Wolfram Mathematica 12.1 selecting as initial parameters the values reported in the literature for hepatocytes (see Supporting Information Table S2). The results of the strain nucleus simulation due to actomyosin contractility as function of ECM stiffness are presented in Figure 3E, where it is observed that the nuclear strain increases when the *k*_*l*_ element behaves like a glassy solid close to its glass transition temperature. This condition represents the dynamics of the cytoskeletal filament network that surrounds the cell nucleus, in contrast with the typical constant value found for a linear elastic model, which in contrast presents no time dependence. All this suggests that phenomena such as nuclear softening^115^ and stiffening^11,123^ are not only regulated by the intrinsic structural components of the nucleus but also by the network of filaments surrounding it and conferring some structural support to the cell nucleus. In Figure 3F, the parameters associated with the shape of the nucleus are presented. With them, it was possible to calculate the strain, shape factor, aspect ratio and Poisson ratio in any specific instant (please see experimental section).

### Nuclear mechanical behavior of hepatocytes

#### Shape evolution of primary hepatocytes

To elucidate the influence of substrate stiffness on the nuclear shape, area, height, deformation and Poisson ratio, the nuclei of primary hepatocytes from Wistar rats plated on elastic polyacrylamide hydrogels were measured at different culture times of 2, 36, 72 and 120 hours. The gels stiffnesses of 1 kPa (physiologically soft) and 23 kPa (stiff) were selected to mimic the natural rigidities of a healthy and fibrotic liver respectively^51,122,124^. In **Figure 4A** and 4B, representative micrographs showing nuclei marked with DAPI as well as CK-18 and F-actin together with the merged channels are presented for soft and stiff respectively. Data of area, height, fluorescence intensity, shape factor and aspect ratio were obtained with a homemade segmentation program written in Python 3.0 and then processed and analyzed with a Wolfram Mathematica 12.1 notebook (see experimental section and supporting information for more details on such tools) (NOTE FOR REVIEWERS: this will be available in a Github repository or upon request at the time of publication). Histograms as a function of their probability density function (PDF) were finally obtained for each condition; more than 200 nuclei were analyzed.

**Figure 4.**
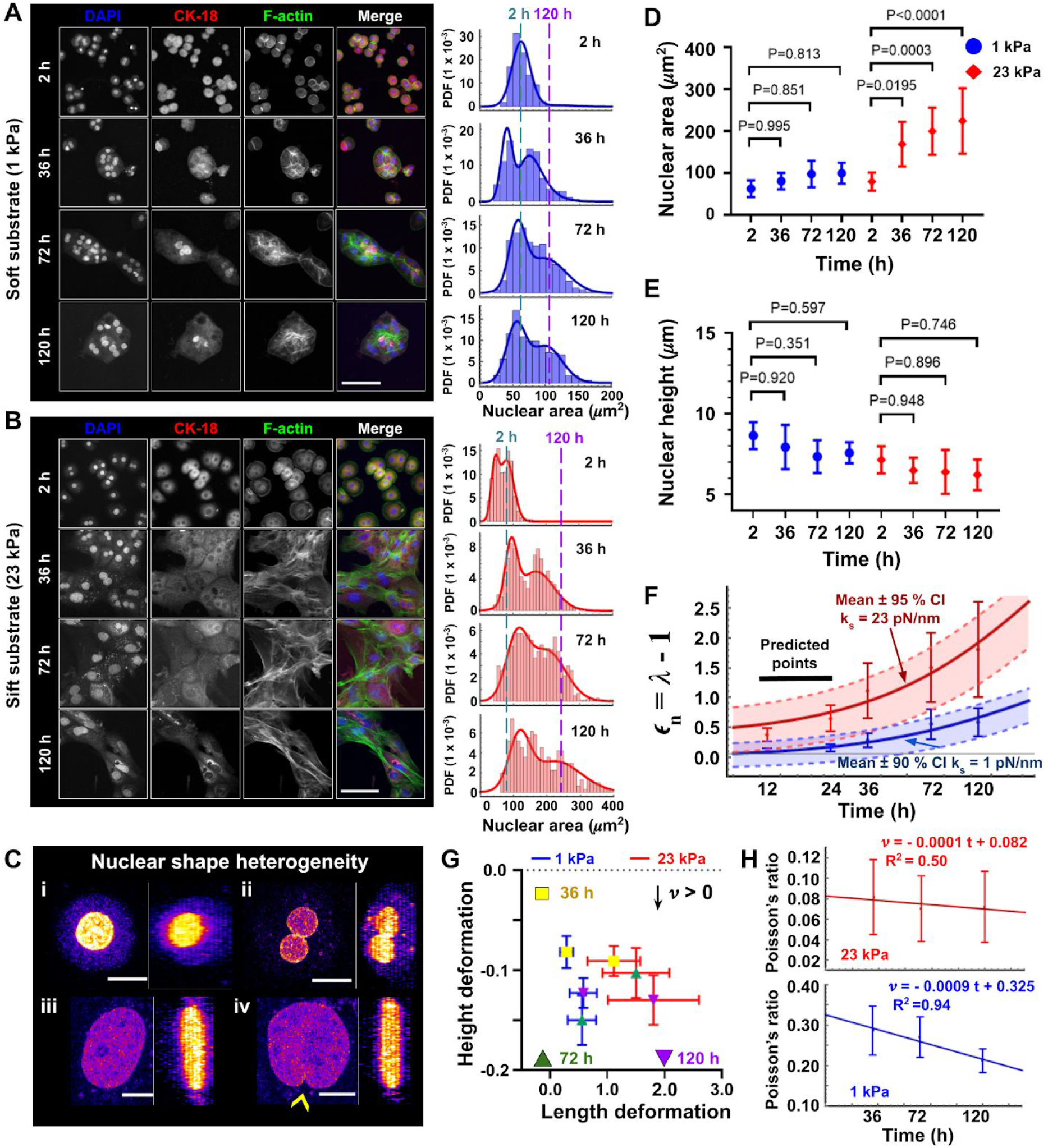
Evolution of the nuclear shape of fresh primary hepatocytes. A) Representative micrographs of fresh hepatocytes seeded on polyacrylamide hydrogels (HG) with a 1 kPa stiffness and conjugated with type I collagen. Cells were cultured and fixed at 2, 36, 72 y 120 hours and the DAPI (blue), CK-18 (red), F-actin (green) channels are shown, together with the merge of all three channels. B) Same results as panel A but in the stiff condition of a 23 kPa hydrogel. In both A and B, the right-hand side histograms depict the projected areas of the nuclei as a function of their probability density function (PDF). The deconvolution of the associated PDF reflects the existence of two sizes; while one is maintained practically constant, the second increases its projected area with time. The vertical dashed lines show the maximal change of projected area occurred between 2h and 120h of culture. C) Representative nuclear shapes observed in the experiments, showing (i) mononuclear and (ii) binuclear cells, that may explain the two PDF populations found in the histograms. The nuclei presented in (iii) and (iv) both belong to the 23 kPa stiff condition after 120h, but their different morphologies may originate from a small indentation, depicted by a yellow arrow in (iv) and a probable consequence of microtubules on a soft nucleus. Panels D) and E) show the average ± SD of the area and height, respectively, of the cell population suffering the greatest change of their projected area. Only a significant difference is observed for the area of hepatocytes cultured on 23 kPa HG, compared to their control of 2h. F) H-VM experimental data and fitting of the nuclear strain at 36, 72 and 120 hours of culture together with a prediction (with a confidence interval of at least 90%) of the non-measured deformation at 12 and 24 hours (see Supporting Information S8 for more details). G) Longitudinal deformation (in the plane of the projected area) vs. the height deformation (orthogonal to the plane of the projected area). Both transverse and longitudinal deformation vary continuously in the two conditions, showing that the deformation is time dependent for a long time (yellow squares = 36h, green triangle = 72h and purple triangle = 120h). From these data, clearly different for the two conditions of soft and stiff HG, the Poisson ratio was calculated. H) Decrease of Poisson ratio in time is observed for the nuclei, and a linear fitting suggests an auxetic behavior. Scale bars: 60 μm in A), 80 μm in B), 10 μm in C). Data presented in D) and E) were analyzed with a one-way analysis of variance (one way ANOVA) with a Tukey multiple comparison correction, results were considered statistically significant when *p* < 0.05.

All the histograms associated with the properties of interest of the nuclei of the primary hepatocytes are shown in the Supporting Information (Figures S4 and S5). Interestingly, two distinct populations of nuclear sizes were found in both stiffness conditions. In the case of the soft substrate, one population maintained its area almost constant (Figure 4A and S4) while the other increased by almost 40% after 120 hours in culture. This behavior is even strengthened on the stiff substrate for which both populations increased in size; one by approximately 300% (from 79.91 ± 21.58 μm^2^ at 2 h to 224.53 ± 78.13 μm^2^ at 120 h) and the other one similarly to what was observed for 1 kPa after 120 hours in culture (in soft gels the largest projected area at 120 h was 99.80 ± 24.87 μm^2^ while in the stiff condition the lowest one was 116.88 ± 30.54 μm^2^ at 120 h, see Figure S4 and S5). The existence of two subpopulation sizes may be related to the nature of the hepatocytes: some are mononuclear cells while others are binuclear, confirmed in our experiments as seen in Figure 4C-i and 4C-ii. ImageJ was then used to establish which of the two populations was the one that may be adequately approximated by the cell mechanical representation shown in Figure 3B and corresponding to cell spreading due to ECM stiffness. The results of the manual quantification of the projected area, shape factor and eccentricity of the nuclei is presented in Figure S9 of the Supporting Information. Figures 4D and 4E present the temporal evolution of the projected area and height of primary hepatocytes cultured on both stiffness conditions. The reference condition is represented here by the cells after 2h of culture, as it is considered that it corresponds to a time when adhesion is guaranteed but before any internal modification happens and with the cells in a stable state^116^. Panel D of that figure depicts clearly that the nuclei of cells cultured on the stiff substrate suffer a greater deformation than the ones on the soft condition, which are almost not deformed at all and remain constant. Moreover, although it is not significant, a decrease in the height of the nuclei is observed in panel D.

Given that the robustness of the generalized Maxwell model (on which the H-VM is principally based here) depends strongly on the obtention of its coefficients from the fitting of experimental data, the comparison between experimental data and the proposed model is presented in Figure 4F. The actual measured data were used to harvest the following constant parameters and coefficients: *k*_*s*0_ (stiffness of the substrate in the reference condition of 2h culture as said above), *k*_*l*0_ (long-term stiffness of the reference condition), *a, b* and *x*_*eff*_ (or *a’, b’* and *c’*). This condition was the one used for the calibration of the H-VM model and to calculate the Cauchy strain (Equation 12) as well as the strain values at 36, 72 and 120 h in both conditions. The reader is invited to revise Supporting Information S1 for a detailed description of the theoretical calculation of the strain and a more detailed explanation of the calibration process. To calculate the cellular nuclear strain from the projected area, the following relation was used:

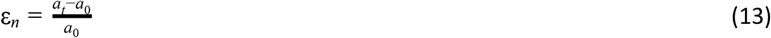

where *a*_*t*_ is the projected area at a time *t* and *a*_*0*_ is the projected area of a reference state. We considered that the projected area is a consequence of the tension generated by the actomyosin apparatus as a function of ECM stiffness, since this tension is on the same plane as that of the projected area^10,125,126^. It is thus valid to compare the nuclear strain obtained from experimental data using Equation 13 and the H-VM model results. Importantly, it was found that the experimental tendency of *k*_*l*_(*t*) better fits a glassy dynamics behavior (see Figure 4F) and does not remain constant over time (thus confirming it is not a linear elastic or hyperelastic material see Figure S1B).

The behavior of *k*_*l*_ (t) was determined by fitting the experimental values of the nuclear strain at different times and on both stiffness conditions with our model (in Wolfram Mathematica 12.1). Equations 10.1 and 10.2 were tested and the values of the R-squared and Akaike information criterion (AIC)^127^ parameters were used to define the most adequate equation for *k*_*l*_(t) (see Table S3). In these experiments, the weak power law of Equation 10.1 did fit better (see Supporting Information S1), inferring that the long-term response of the nuclei of the primary cells follow a glassy dynamic behavior at long timescales. Then, the now complete model calibrated with the experimental data was successfully tested for its capability to extract information of mechanical configurations prior to the measured (and adjusted) data of 36, 72 and 120 hours. The nuclear strain extrapolated at 12 and 24 hours was then validated with experimental results obtained from two additional independent experiments with fresh primary hepatocytes cultured on the same soft and stiff conditions but fixed and stained at 12h and 24h (results are presented in Figure S10 of the Supporting Information). Figure 4F shows that the H-VM construct is capable of revealing previous strain values from the outcome of these experiments in primary liver cells for shorter times of culture. For the soft hydrogel, the strain values predicted by the model at 12 and 24 h with a 90 % of confidence interval are 0.09 ± 0.18 and 0.17 ± 0.19 respectively which is very close to the experimentals results: 0.1 ± 0.04 at 12 h and 0.16 ± 0.06 at 24 h. For the sift conditions the strain values predicted by the model at 12 and 24 h with a 95 % of confidence interval are: 0.56 ± 0.37 and 0.75 ± 0.38 respectively, close to the experimental data were 0.38 ± 0.10 and 0.65 ± 0.22.

Interestingly, it is also possible to compute the stiffness *k*_*s*0_ of the last stable reference condition corresponding to the healthy liver from which the cells came from before culture on a different material. In this case, the model allocated a *k*_*s*0_ value of 0.75 pN/nm, in very close agreement with the literature^118,128^ where it is reported that the stiffness of the hepatocytes ranges between 0.75 to 1 kPa. Using the Boussinesq Green’s function approach^129^ and a length scale of 1 *μ*m^126^, this is equivalent to approximately 0.75 - 1 pN/nm. The stiffness of the healthy liver is reported to be ~1.12 ± 0.15 which is equivalent to ~1.12 ± 0.15 pN/nm (see Supporting Information S1 and Table S1) From the dynamics of the H-VM model, it is possible to predict that at long timescales the nucleus of primary hepatocytes will probably acquire a highly dissipative state on such substrates with a stiffness *k*_*s*_ ≻ *k*_*s*0_ (*k*_*s*0_ of a healthy liver in this case). Indeed, in these conditions, the *k*_*l*_ value associated with the elastic energy storage (according to Equation 10.1) will tend to decrease with time while τ _*NE*_ (hence the NE viscosity) will increase. This behavior may be explored in detail by the reader, using the computed application mentioned above.

Finally, Figure 4G presents the graph of the deformation of the height *h* as a function of the longitudinal strain associated with the deformation calculated with the Equation 13. This experimental result shows that the deformation is time dependent over a long time, since it varies continuously in the two stiffness conditions. This experimental data also helped determine the Poisson ratio for both stiffness conditions and over time, as presented in Figure 4H. As a first approximation, the average was calculated over all the time points obtaining ν ≈ 0.074 ± 0.006 and ν ≈ 0.255 ± 0.039 for the stiff and soft condition respectively. However, as observed in panel 4H, the Poisson’s ratio presents a time-dependent behavior, as previously observed in the literature for living cells^130^. A linear fitting was also performed showing a good fit for the soft condition (*R*^2^ = 0.94), although it was not acceptable for the stiff condition (*R*^2^ = 0.50). Interestingly, an automated best fit provided by the software revealed a nonlinear behavior of the Poisson ratio of the nucleus, for the stift condition the best fit was 0.104 − 7.77 × 10^−4^*t* + 4.26 × 10^−6^*t*^2^ with a correlation coefficient of *R*^2^ = 0.999 and for the soft condition it was 0.27 + 7.98 × 10^−4^*t* − 1.07 × 10^−5^*t*^2^ with a correlation coefficient of *R*^2^ = 0.999. The change in sign of the linear coefficient suggests that the nuclear volume has two phases in time and this may be dependent on the stiffness of the substrate on which the cell is deposited). Additionally, the decrease observed in the Poisson’s ratio at long times presented in Figure 4H is also representative of materials that exhibit a glassy behavior^130^. In addition to this, to verify whether the nucleus of cells fixed at 2 hours of culture is a good reference point, the logarithmic strain was used to calculate the pre-strained of the nucleus. A value of 0.993 was found (see Supporting information S1 and figure S10 panel E), meaning that the nucleus of the primary hepatocytes presents a prestress of 0.397 kPa, which is of the same order of magnitude as the common value of 0.5 kPa^131,132^.

### Shape and area evolution of HepG2 cell line

A human hepatocellular carcinoma cell line was then cultured and studied in the same conditions for comparison. Indeed, it was assumed that the reference stable mechanical condition of these HepG2 cells would differ from that of the primary hepatocytes described above, due to an evidently different mechanical behavior originating from an altered mechanical state of a cancer even though they still preserve epithelial characteristics and respond to stiffness^133^.

The results of the temporal evolution of the nuclear shape for HepG2 cells is presented in **Figure 5**. Panels A and B show the representative micrographs associated with the different stiffness conditions and it may be observed that two dynamic populations were found in the histograms, similarly to primary hepatocytes. However, in this case the maximal change in projected area occurred at 72h and decreased afterwards, while it happened after at least 120h of culture for primary hepatocytes. This is more evident in panels C and D where a reduction in the projected area is seen, as well as an increase in height of the nucleus in the soft condition. This tendency is even clearer in panel E where the nonlinear behavior of *ϵ*(*t*) is noticeable. In this case, the fitting of the evolution of the nuclear deformation using Equations 10.1 and 10.2 led to a different behavior: *k*_*l*_(*t*) = *a*′ − *b*′*t* + *c*′ *t*^2^. In the context of glassy dynamics, this implies that *k*_*l*_(*t*) does not acquire a stable state over time for HepG2. This could be associated with an aging-like phenomenon, because the subcomponents behind *k*_*l*_(*t*) (it was assumed here that they are the filaments around the nucleus) are not capable of accessing all possible energetic states, since the material presents a really slow recovery of their rheological properties. This, in turn, indicates that *P*(*E*_*T*_, *l*|*t*) does not present a constant behavior in time. Interestingly, in soft matter theory this statement is precisely the definition of aging^67^. Additionally, in panel E it is observed that the strain suffered by the nucleus of the cell in the soft condition before 72 h is greater than the strain of the nucleus of the line in the stift condition.

**Figure 5.**
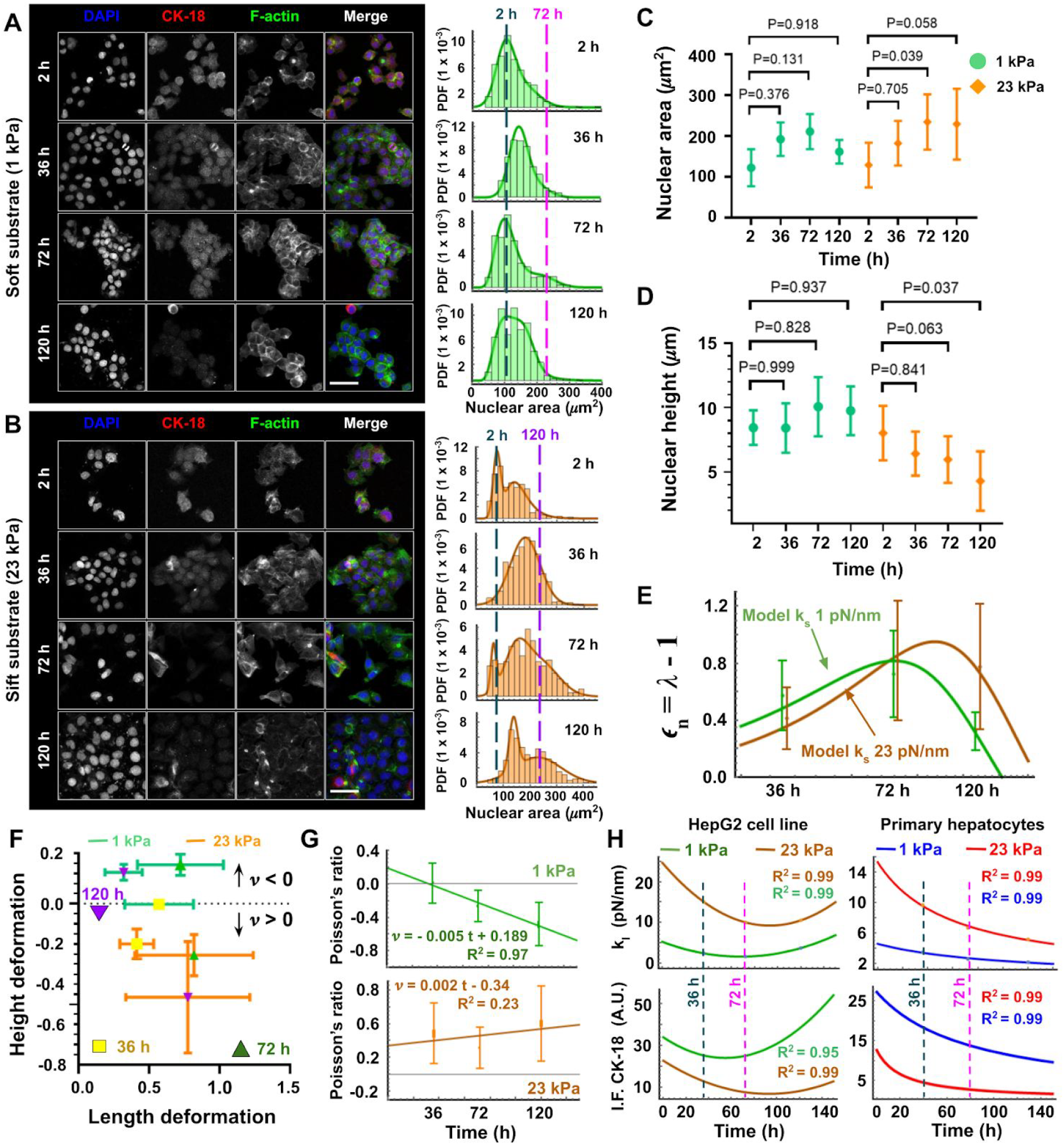
Evolution of nuclear shape of HepG2 cell line. The results are presented similarly to those of Figure 4, in order to compare the nuclear behaviour of a primary cell (originating *a priori* from a mechanically stable state of homeostasis condition) with a cancerous cell line (*a priori* starting far from the natural mechanical state of a soft liver). A) Representative micrographs of HepG2 cells seeded on polyacrylamide hydrogels (HG) with a 1 kPa stiffness and conjugated with type I collagen. Cells were cultured and fixed at 2, 36, 72 y 120 hours and the DAPI (blue), CK-18 (red), F-actin (green) channels are shown, together with the merge of all three channels. B) Same results as panel A but in the stiff condition of a 23 kPa hydrogel. In both A and B, the right-hand side histograms still depict the projected areas of the nuclei as a function of their probability density function (PDF). The deconvolution of the associated PDF reflects the existence of two sizes; while one is maintained practically constant, the second increases its projected area with time. The vertical dashed lines show the maximal change of projected area occurred between 2h and 72h of culture for soft substrates and between 2h and 120h for stiff ones. Panels C) and D) show the average ± SD of the area and height, respectively, of the cell population suffering the greatest change of their projected area. Once again, only a significant difference is observed for the area of HepG2 cultured at 23 kPa HG. E) Comparison between experimental data of panel C and the H-VM fitting of the nuclear strain using Equation 13. F) Longitudinal deformation (in the plane of the projected area) vs. the height deformation (orthogonal to the plane of the projected area). The longitudinal deformation is greater before the 72 hours in culture and then the absolute height deformation increases, differently to what was observed for fresh primary cells where two continuous deformations were observed for the whole duration of the experiments. Moreover, here the substrate stiffness seems important in the cell behavior. G) Decreasing of Poisson ratio in time is observed for the HepG2 nuclei on soft substrates, and a linear fitting suggests an auxetic behavior of the nuclei on both conditions after 72h. H) Comparison between the fitting functions of *k*_*l*_(*t*) for the data from HepG2 cells and primary hepatocytes on two different stiffnesses (top left and top right); and comparison between the intensity of fluorescence of CK-18 for both cells (bottom). Scale bars: 60 μm in A), 60 μm in B). Data presented in C) and D) were analyzed with a one-way analysis of variance (one way ANOVA) with a Tukey multiple comparison correction, results were considered statistically significant when *p* < 0.05.

These results pointing towards an aging process for HepG2 cell line are also reinforced by the measurement of the nuclear Poisson ratio (Figure 5G): it presents a linear dependence with time, a phenomenon that has been associated with processes of cell cytoskeleton reorganization^115^ and aging^134^. Moreover, when analyzing the longitudinal and height deformations presented in Figure 5F, it was very interesting to find a clear auxetic behavior of the nucleus of this cancer cell line on the soft condition. This uncommon mechanical response has already been observed in non-differentiated stem cells^135^, where Pagliara and collaborators assumed that the auxetic behavior of the nucleus was a consequence of the cells undergoing a transitional metastable state through which they acquire a differentiated state irreversibly. A similar metastable state, that can then be related to a certain form of mechanical aging, is actually observed here in the behavior of *k*_*l*_(*t*) presented in Figure 2C and 5H for *x*_*eff*_ < *x*_*g*_.

Finally, to test our hypothesis on the nature of the subcellular components governing the behavior of the *k*_*l*_(*t*) mechanical element, the intensity of fluorescence (IF) of CK-18 was quantified for both cell types (see Supporting Information Figures S10.1 and 10.2). Figure 5H shows the comparisons between the two cells types when fitting experimental data of both *k*_*l*_(*t*) and IF(CK-18) on the two different stiffnesses. Interestingly, *k*_*l*_(*t*) and IF(CK-18) presented very similar temporal evolutions for each cell type, but HepG2 and primary hepatocytes behave differently. Primary hepatocytes clearly show a decrease in CK-18 intensity over time, a tendency that was also well fitted by a weak power law (Equation 10.1) but once again the tendency of HepG2 cells resulted better described by Equation 10.2. Interestingly, it has been reported in the literature that the half life of such cytokeratins present in liver cells is between 84 and 120 hours^121^. This may then explain why the nuclear shape is significantly modified around this time when in culture. Moreover, the difference in the magnitude of IF(CK-18) between soft and stiff conditions may also suggest that CK-18 is not the only intermediate filament involved in the irreversibility process.

## Discussion

### Glassy dynamics is an accurate model of the long-term response of cell nucleus

One of the main mechanical functions of the nucleus is to maintain the spatial organization that controls the openness and repression of its sequences, and the structural integrity of the genome. Its active response to mechanical stimuli is being studied and modeled in several research works, together with the individual and collaborative contribution of all the structural components that govern its shape. One of the main goals is to better understand the underlying mechanisms that modify its mechanical properties to guarantee a stable and functional configuration during homeostasis or in response to intra- and extracellular mechanical stimuli. The relevance of the H-VM lies in the fact that it not only describes and emulates the viscoelastic response of the nuclear envelope and chromatin but it also integrates their active response to mechanical perturbations, based on experimental observations. In this context, it is then proposed that the long time behavior of cell nucleus (hours or days) is dictated not only by the active behavior of the NE, but also by the dynamic reorganization of the filaments network surrounding it. Thanks to its calibration and validation with experimental data, we demonstrate that the proposed model is suitable for modeling of the nucleus mechanical behavior at both short and long timescales.

It is clear in the simulations of the H-VM model (see Figure S1B and S1C) that by only modeling the nucleus as a linear viscoelastic material (Figure 1B) and incorporating the active behavior of its NE (Figure 2A), the short-term mechanical response of the nucleus is correctly depicted (see Figure 1C, 1D and 1E). However, the long-term nuclear response, that is changing in time (Figure 4F and 5E), is not strictly respected. An important contribution of the model was thus to assume a glassy dynamics behavior for the nucleus and an active response of the NE that determine the long-term dynamics of the nuclear mechanical behavior. Additionally, it is demonstrated that by calibrating the parameters of the model with experimental data, it seems possible to determine the characteristics of the last stable mechanical state sensed by the nucleus, including the parameters governing the glassy nuclear dynamics (using Equations 10.1 and 10.2). Finally, it may be inferred that this reference state is specific to each cell type, probably dependent on its origin and prior mechanical conditions as a result of their spatial organization and mechanical properties of their particular filaments network.

### The nucleus projected area is proportional to substrate stiffness, but deformation is not

By analyzing the temporal evolution of the projected area of nuclei belonging to primary hepatocytes and human hepatocellular carcinoma cell line HepG2 at different times of culture, it is found that the average projected area of the nuclei of the primary cells is smaller than that of the cancer cell line (see Figure S4 to S7). This is somehow expected, as it was reported that the nuclei of cancer cells are usually greater than normal cells^136,137^. Besides, it was observed in particular that the overall deformation was larger for the primary cells than for the cell line and that the dynamics of deformation also differed (Figures 4F and 5E). On one hand, primary cells increased their nucleus size gradually, at least until 120 h of culture, with the substrate stiffness intensifying this effect (Figure 4F). On the other hand, HepG2 presented a maximal deformation after 72 h and in this case the deformation was larger on the softest substrate before 72 h (Figure 5E). To explain this, the H-VM suggests an explanation of what may happen if the nucleus is under a mechanical tension different from that of the cell reference state: the expression levels of lamin A and lamin B1 increase if the ECM stiffness became greater than that of the physiological reference state or they decrease if the ECM became softer. This effect then leads to a respectively higher or lower deformability of the nucleus as a function of the substrate stiffness, in collaboration with the glassy dynamics associated with the network of filaments surrounding the nuclear envelope. Therefore, it may be assumed that the twofold deformation of the primary cells nuclei on hydrogels substrates, compared to the cell line, is due to a greater capacity of the cell line nucleus to dissipate elastic energy. This higher capacity of dissipation probably originates from a stiffer natural environment (reference state) for the cancer cell line, causing lower deformation on soft gels. These observations confirm that despite a dependency of the projected area on the substrate stiffness (Figures 4D and 5C), the actual deformation depends more on the cell type and the reference mechanical state of its nucleus (Figures 4F and 5E).

### The nucleus shape may store information of previous mechanical states

In the continuous mechanics theory, a mechanical deformation is defined as the transformation of a body from a previous reference configuration (a stable *preconfiguration* referring to a specific spatial organization of all particles that compose the object in question) towards a current configuration^138^. If the deformation can be described by a linear transformation that preserves lines and parallelism, it is called an affine deformation^139^ and in this case all the geometrical information of the body may be described by a dimensionless parameter called strain which is a consequence of the physical properties of the body and the sum of stresses exerted on it. In this context arose the hypothesis that a current nuclear shape stored information from previous mechanical states, despite the fact that most of the mechanotransduction processes occur in short periods of time^1^. Since the nucleus is not an elastic material and presents an irreversible mechanical behavior, its shape may be considered as the result of the history of forces that deformed it. And this storage of the mechanical information may then be revealed by adjusting the H-VM to the nuclear strain obtained experimentally.

With this approach, by calibrating the model with experimental results at times of 36, 72 and 120 hours (see Supporting Information S1), the model was capable of revealing the magnitude of the nuclear deformation corresponding to previous states that were not used to calibrate the model (Figure 4F). Indeed, the values of the magnitude of deformation calculated for shorter times of 12 and 24 hours matched very well with the experimental results of a different set of independent experiments with other primary hepatocytes. Indeed, the predictions fell within a 95% confidence interval for nuclei of cells in the stiff condition and with 90% in the soft condition (Figure 4F).

Regarding the reference mechanical state, specific to a cell (referring to a previous stable state before perturbation), the model resulted very useful for the determination of its mechanical properties. For fresh primary hepatocytes, the original basal stiffness *k*_*s*0_ (liver tissue) was found at 0.75 pN/nm with a long-term stiffness *k*_*l*0_ of 4 pN/nm and a nuclear envelope relaxation time of τ _*NE*_ ≈ 1 *s*. It is important to mention that these values are in very close agreement with a healthy liver^118^. In contrast, for HepG2 (a modified cancer cell line optimized for *in vitro* culture on stiff substrates coated with collagen I), the reference stable state indicated a much greater basal stiffness *k*_*s*0_ of 8 pN/nm with a long-term stiffness *k*_*l*0_ of 15 pN/nm and a much longer nuclear relaxation time τ _*NE*_ ≈ 25.1 *s*. Noticeably, the *k*_*s*0_ value of 8 pN/nm ~ 8 kPa for the stiffness of the original substrate of HepG2 (reference state) seems very high for a liver tissue, but is much lower than a common culture plate for which this cell line has been selected and is viable, providing an adhesion protein like collagen I is coating it. This may suggest that the *k*_*s*0_ here rather reflects the external stiffness sensed by the nucleus (and to which it responds), and this is probably why it is different from the GPa range of plastics or glass used as the contractile force generated by the actomyosin apparatus is reported to tend to a plateau because it is proportional to cell spreading^10^, thus limiting the tension supported by the nucleus. In the context of this observation, it is also important to notice that cancer cells lose some capacity to sense and respond to substrate stiffness^140^, which may also explain the value found here. Finally, another contribution of this work is to suggest that it is necessary to monitor subcellular proteins constituting the network surrounding the nucleus for long periods of time, at least longer than their reported characteristic half-lives, in order to observe their potential impact on the eventual nuclear shape, or to monitor their post-translational modifications that affect their functions. Indeed, the H-VM indicates that the dynamics responsible for the process of replacement of these proteins, as well as their spatial organization as a network, both play an important role in the nuclear dynamics. Typical times of 70 hours up to 120 hours have been reported for histones and intermediate filaments respectively, corresponding to the apparition of significant nuclear changes^97^.

### Glassy dynamics applied to long timescales unveils a mechanical aging process

Another noticeable difference was that the nuclei of primary hepatocytes and HepG2 exhibited different time behaviors. Primary cells showed a quasilinear behavior described by a weak power law (Equation 10.1). Using soft glass matter theory, this implies that the force perceived by the nucleus is proportional to the temporal change in its deformation, rather than to the deformation itself. Biologically, this can be interpreted by a tendency of the force perceived by the nucleus to reach a stationary state more rapidly than the deformation, i.e. dissipative phenomena and viscosity strongly dominate the nuclear shape. For primary hepatocytes in culture, this resistance seems to be lost when cells start expressing vimentin (or lose CK-18); and that precisely coincides with a deformation of the nucleus^141,142^. In contrast, HepG2 nuclei followed a nonlinear behavior (Equation 10.2) originated by the reduction of the energy states that the nuclear components may access; in other words, the modifications of the mechanical properties of the nucleus has a high energy cost related to aging processes^67,120^. This was explicitly manifested by the calculation of the Poisson’s ratio: the nuclei of primary cells showed a Poisson ratio ranging between 0.1 and 0.4 while HepG2 nuclei demonstrated an auxetic behavior on soft substrates. This suggests that the 1 kPa condition did relax the tension on the cellular nucleus of the cancer cells, being responsible for such a change in the relation between transverse and longitudinal strains. This is of particular importance in cell biology and will need further study as auxetic behaviors have been observed to be related to a certain form of ageing caused by mechanical stress^134^. In this direction, it is also relevant to note that in spite of a similar epithelial origin, the cells used here have different “natural” microenvironments; primary hepatocytes were isolated from a mechanical homeostasis state while HepG2 are originated from an altered state of a human hepatocellular carcinoma and were preserved in stiff polystyrene Petri dishes. Additionally, these findings show that the hypothesis of incompressibility commonly used in the mechanical models of the nucleus (where ν ≈ 0.5) is actually not fulfilled. Furthermore, the temporal behavior of nuclear deformations strongly suggests that it is governed by the reorganization of the cytoskeleton on time scales of the order of hours to days.

## Conclusion

To study the mechanical response of the nucleus at different timescales, a hybrid viscoelastic model integrating both continuum mechanics and soft glass matter theory was developed, in order to integrate the instantaneous viscoelastic response of the structural components of the nucleus as well as the active response of the nuclear envelope and the dynamic reorganization of the cytoskeleton at different timescales. Given that the H-VM is based on a viscoelastic model with 5 elements in the representation of the generalized Maxwell model, its calibration using experimental data is absolutely necessary. However, although the model depends on so many parameters, the only free parameters with which the model is adjusted are *k*_*s*0_, *k*_*l*0_ and *k*_*l*_. Indeed, with these parameters, all the others (that are not fixed by the experimental conditions) are found, thus avoiding an overadjustment of the system. Furthermore, it is shown that in spite of a 1D formulation of the H-VM, it is capable of reproducing the phenomenon of nuclear strain at long term. These results suggest that the mechanical response of the nucleus of liver cells is consistent with the hypothesis that the elements responsible for the long-term mechanical behavior of the nucleus are both the active response of the nuclear envelope and the continuous spatial reorganization of the filament network that surround the nucleus.

Additionally, based on experimental evidence, the H-VM may be used to reveal that the increase in the nuclear strain at long timescales implies a nuclear softening, a phenomenon accelerated if the stiffness of the cell substrate is higher. However, this response is accompanied by an increase of the dissipative properties of the nucleus, granting it a certain structural stability. Moreover, it is observed that cell types adapted to stiff substrates and originated from an altered mechanical state, such as HepG2 hepatocellular carcinoma cells, manifest a temporal behavior more complex and that is even exacerbated on soft substrates and longer timescales. In this context, the H-VM was able to reveal processes of aging and mechanical memory, which are consistent with the dynamic behavior of the Poisson’s ratio of the nuclei measured experimentally Finally, this work demonstrates the need to use both continuum mechanics and mesoscopic soft matter theory to describe the dissipative phenomena of the nucleus. It is believed that this work will be helpful to cell biologists and biophysicists who need to model both short and long-term nuclear mechanical behaviors to explain experimental results, offering a broader view of the distinct nuclear responses to the extracellular mechanics.

## Acknowledgements

The authors would like to thank DGAPA-PAPIIT IT102017 for financial support and LaNSBioDyT National Laboratory where the research was performed. DPC is a doctoral student from the Programa de Doctorado en Ciencia e Ingeniería de Materiales, Universidad Nacional Autónoma de México (UNAM) and has received CONACyT fellowship 594952. The authors acknowledge Adriana Rodríguez-Hernández warmly for her kind help with the cell culture.

