## Supplementary material and figures for "Elucidating the short and long-term mechanical response of the cell nucleus with a hybrid-viscoelastic model"

### Electronic supplementary information (ESI)

[b]*Departamento de Física*
*Facultad de Ciencias, Universidad Nacional Autónoma de México*
*Circuito Exterior S/N, Ciudad Universitaria CP 04510, Coyoacán, CDMX, México*

[c]*Posgrado en Ciencia e Ingeniería de Materiales*
*Facultad de Ciencias, Universidad Nacional Autónoma de México*
*Circuito Exterior S/N, Ciudad Universitaria CP 04510, Coyoacán, CDMX, México*

[d]*Posgrado en Ingeniería*
*Facultad de Ciencias, Universidad Nacional Autónoma de México*
*Circuito Exterior S/N, Ciudad Universitaria CP 04510, Coyoacán, CDMX, México*

[e]*Unidad de Imagenología Cuantitativa*
*Facultad de Ciencias, Universidad Nacional Autónoma de México*
*Circuito Exterior S/N, Ciudad Universitaria CP 04510, Coyoacán, CDMX, México*

#### S1. Representation of the mechanical elements of the cytoskeleton and adhesion complex

The complete representation of all the mechanical elements of an adherent cell (nucleus-cytoskeleton-adhesion) was based on the union of the H-VM nuclear model with the mechanical elements reported by Cao and collaborators[1] (see Figure 4B). The constitutive equations in one dimension (1D) that represent these mechanical elements are presented below:

$$F_a = k_a\, x_a \tag{S1}$$

$$F_c = \varrho + k\, x_{\mu T} \tag{S2}$$

with

$$\varrho = \frac{\beta\, \rho_0}{(\beta - \alpha)} + \frac{\alpha\, k_{\mu T} - 1}{\beta - \alpha} x_{\mu T} \tag{S3}$$

Also

$$F_{FA} = k_{eff}\, x_{FA} \tag{S4}$$

with

$$k_{eff} = \frac{\sqrt{k_p k_s k_c}\ (k_p + k_s)^{3/2}\ sinh(L/L_c)}{(k_p^2 + k_s^2)\ cosh(L/L_c) + k_p k_s\ (\ 2 + (L/L_c)\ sinh(L/L_c)} \tag{S5}$$

$$L_C = d_c \sqrt{\frac{1}{k_c} \frac{1}{(1/k_s + 1/k_c)}} \tag{S6}$$

Equation S1 represents the force $F_a$ supported by the actin filaments with a stiffness $k_a$ exerted by the contractility $F_c$ of the actomyosin apparatus. Equations S2 and S3 describe an active contractile element with positive feedback: $\varrho_0$ is the basal contractility of the cell in the absence of external stress or restriction (it may be interpreted as the intrinsic contractile tension, or cell pre-stress). The parameters $\alpha$ and [illegible] represent the mechanochemical coupling parameters respectively and are associated with a molecular mechanism reflecting the stress-dependent signaling pathways and engagement of motors and they satisfy the conditions $0 < \alpha/\beta < 1$. Also, $k_{\mu T}$ represents the stiffness of microtubules, hence, $\varrho$ and $k_{\mu T}$ represent the

chemomechanical activity of the myosin motors. Equations S4-S6 describe the mechanical response of the focal adhesion complex (FA) caused by the force transmitted to the actomyosin apparatus and is called the effective stiffness of the FA and ECM (FA- ECM). where $L_C$ is the shear lag length, which is defined as the length scale where actomyosin forces are transmitted. $ks$ , $k_p$ and $k_c$ are the stiffnesses associated with the substrate/ECM, the adhesion plaque and the clutch, respectively. $d_c$ is the spacing between integrins and *L* is the focal adhesion length.
Finally, $x_a$, $x_{\mu T}$ and $x_{FA}$ are the deformations of the actin filaments, of the microtubules and the FA respectively. These equations are detailed precisely in the work of Shenoy group[2,3].

In order to calculate the nuclear strain due to physiological forces using a notebook in Wolfram Mathematica 12.1 (shared upon request and with which the reader can play to visually understand the role of each parameter), the system composed of the Equations 1-3, 8-10 and S1-S6 was solved following the procedure and assumptions described below:

1. As the components of this model are connected in series, we have:

$$F_n = F_a = F_c = F_{FA} \quad \text{(S7)}$$
$$x = x_n + x_a + x_c + x_{FA} \quad \text{(S8)}$$

2. Since we assume that the cell is in a stable state during a short time interval (i.e. $dx/dt = C0 \Rightarrow s\,\overline{x} = C0$ , which implies that the cell does not change its size or if it does, it is at a constant rate), then it is possible to impose a geometrical constraint in the Laplace space:

$$\overline{x} = \overline{x_n} + \overline{x_a} + \overline{x_c} + \overline{x_{FA}} = C0/s \quad \text{(S-L1)}$$

where *C0* is a constant, for simplicity C0 was set to 0 since at long term, the spreading of the cell reaches a stable value (constant).

3. For simplicity, Equations 1-3 and S1-S6 were transformed to the Laplace space assuming constant parameters and $F_i(t) \wedge x_i(t)$, then, using the chain rule in the Equations 1-9 we derived with respect to time and obtained:

$$\overline{F_n} = \overline{F} = k_l\overline{x_n} + \sum_j^2 \frac{k_j\,\overline{x_n}}{1+1/s\,\tau_j} \quad \text{(S-L2)}$$
$$\overline{F_a} = \overline{F} = k_a\overline{x_a} \quad \text{(S-L3)}$$
$$\overline{F_c} = \overline{F} = k_{\mu T}\overline{x_{\mu T}} + \frac{\beta\,\rho_0}{(\beta-\alpha)} + \frac{\alpha\,k_{\mu T} - 1}{\beta-\alpha}\overline{x_{\mu T}} \quad \text{(S-L4)}$$
$$\overline{F_{FA}} = \overline{F} = k_{eff}\overline{x_{FA}} \quad \text{(S-L5)}$$

We used the notation $L(F(t)) = \overline{F}$ where *L* is the Laplace transform operator.

4. Then, Equations S-L1 to S-L5 were solved algebraically as a function of $\overline{F(s)}$ in the Laplace space.

5. In order to find the nuclear strain, the Boltzmann equation was used (Equation 11 in the main text) with $\overline{\varepsilon(s)}$ and the creep modulus $\overline{J(s)}$ of the cell nucleus in the Laplace space, leading to:

$$\overline{J(s)} = F_0/(k_l + \sum_j^2 \frac{k_j}{1+1/s\,\tau_j}) \quad \text{(S-L7)}$$
$$\overline{\varepsilon(s)} = \overline{J(s)} * \overline{F(s)} \quad \text{(S-L8)}$$

6. Since it has been shown experimentally that the cell nucleus is under certain stress over time (nuclear pre-stress)[4], mathematically this implies that $\varepsilon(t=0) \neq 0$ because $F(t=0) \neq 0$ . Moreover, Equation S7 establishes that $\varepsilon(0)=0 \Leftrightarrow F(0)=0$, then, to be consistent with the experimental information, it is necessary to assume a reference stable state $\varepsilon(\tau_{NE0}, k_{l0}|k_{s0}) = l_0 \neq 0$ that depends on the substrate/ECM stiffness (we show in the main text Figure 1 and in the Wolfram applications that $\tau_{NE}$ and $k_{l0}$ depend on the ECM stiffness). Then, the relative deformation of the cell nucleus takes the following form:

$$\varepsilon_n(t) = \lambda - 1 = \frac{\varepsilon_n(t) - \varepsilon_{n0}(t_0)}{\varepsilon_{n0}(t_0)} \qquad \text{(S9)}$$

where $\lambda = l/l_0$ is the common definition of strain and $\varepsilon_n(t)$ is the Cauchy strain definition.

7. In order to solve Equation S9 from Equation S-L8 we used the inverse Laplace transform $L^{-1}$. However, as shown in the main text, $\tau_{NE}$ and $k_l$ possess a dynamic behavior varying in time. Nevertheless, since *F* was calculated from the assumption that the cell is in a stable state, then it is valid to assume that, in a sufficiently short period of time, the value of parameters $\eta_{LA}$, $k_{LB}$ and $k_l$ remain constant. Hence, the strain function of the nucleus is obtained from the extrapolation of those specific time points $t_i$ calculated with the values of $\eta_{LA}$, $k_{LB}$ and $k_l$ evaluated at the time $t_i$. This assumption is valid since the nucleus exhibits continuous mechanical deformation.

8. Finally, due to the nature of the H-VM, which is based on the Generalized Maxwell model, it is necessary to calibrate with actual data and fit them in order to find the values of the parameters $k_{l0}$, $k_{s0}$, b and $x_{eff}$ (or b' and c'). It is important to emphasize that these parameters characterize the reference nuclear configuration. And then, it is possible to obtain the initial (reference) configuration of the nucleus through fitting to at least three experimental points for each stiffness condition. The procedure to calibrate the model is described below:
    a. Using the Wolfram Mathematica piece of software, the equations are solved up to point 7, obtaining a relation of the form $\overline{\varepsilon_i} = \overline{\varepsilon_i}(k_s, k_l, s)$ (S9.1), where the parameters $k_s$ and $k_l$ are free, s in the Laplace parameter and the index i represents each condition, reference state characterized by the values $k_{s0}$ and $k_{l0}$, nucleus in 1 kPa substrate characterized by the values $k_{s1}$ and $k_{l1}$ and nucleus in 23 kPa substrate characterized by the values $k_{s23}$ and $k_{l23}$.
    b. Then, the values $k_{s1}$ and $k_{s23}$ were set to 1 pN/nm and 23 pN/ nm, these equivalence between the elastic modulus and the stiffness was achieve through the Boussinesq Green's function approach[5,6] where $k_s \approx 2rE$ (S9.2), here r is the length scale, since the focal adhesions size is on the order of micrometers[6], then $2r \approx 1\mu m$, so a substrate with a young modulus of 1 kPa has a stiffness of approximately 1 pN/nm.
    c. Then, using the Lapalce inverse transformation on the equation S9.1, the temporal strain equation was obtained. So we fixed the time to t = 129600 s which is equivalent to 36 h and using the equation S9 the system was variaging to fit to the experimental points (Figure 4F and 5E). From this variation of parameters the values $k_{s0}$, $k_{l0}$, $k_{l1}$ and $k_{l23}$ were found at t=36 h, So, the parameters $k_{s0}$ and $k_{l0}$ were fixed.
    d. Finally, with $k_{s0}$ and $k_{l0}$ fixed, the values of $k_{l1}$ and $k_{l23}$ were also found varying these parameters at t= 72 h and t= 120 h. Once the three $k_l$ values were obtained for each stiffness, the equations 10.1 and 10.2 were fitted to them. To determine which one of the two Equations 10 (10.1 or 10.2) best describes the dynamics of $k_l(t)$, the correlation parameter $R^2$ and the Akaike information criterion (AIC)[7], included in the Wolfram Mathematica software, were used. The AIC parameter is widely used to compare the relevance of different models fitted to experimental data.

9. Once the model is calibrated with experimental data, it is possible to predict deformation values within the adjusted time interval and the storage and loss dynamic modules in the frequency space.

$$E'(\omega) = k_l(\omega) + \frac{\omega^2 k_{LB}(\omega)\,\tau_{NE}}{\omega^2\tau_{NE}^2(\omega)-1} + \frac{\omega^2 k_{ch}\,\tau_{ch}}{\omega^2\tau_{ch}^2-1} \quad \text{(S10)}$$

$$E''(\omega) = \frac{\omega\, k_{LB}(\omega)\,\tau_{NE}}{\omega^2\tau_{NE}^2(\omega)-1} + \frac{\omega\, k_{ch}\,\tau_{ch}}{\omega^2\tau_{ch}^2-1} \quad \text{(S11)}$$

10. Additionally using the Henck´s strain or logarithmic strain definition[8], it is possible to calculate the pre-strain of the nucleus.

$$H = dl/L = ln(\lambda) = ln(\tfrac{L}{l_0}\tfrac{l}{L}) = ln(H_0) + ln(H_1) \quad \text{(S12)}$$

where $H_0 \equiv L/l_0$ is the pre-strain of the system and $H_i \equiv l/L$ is the increment of the strain. The distances $l$, $l_0$ and $L$ are the total deformation, the original length and the increment length respectively. It is then possible to apply Equation S12 for two systems with the same pre-strain, since the increment may be assumed as a linear increment. Then, Equation S12 adopts the equation of a line, and we have:

$$ln(\lambda) = mt + b \quad \text{(S13)}$$

where $b$ is the pre-strain. However, since the point $t = 0$ is arbitrary, it is necessary to apply Equation S13 for at least two conditions (in this case, two stiffness conditions) and the interaction among the straight line will be the $ln(H_0) = b_i$.

Finally, since with the H-VM it is possible to obtain the $k_{l0}$ value, then the pre-stress condition may be calculated as follows:

$$F_0 = k_{l0} Exp(b_i) \quad \text{(S14)}$$

Figure S1 depicts some of the results obtained here for primary hepatocytes and HepG2 cells at different times and for soft (1 kPa) and stiff (23 kPa) substrates. On panel A, once the contractility caused by the actomyosin apparatus reaches a constant value over time, the exerted force was calculated with the equations SL1-SL5, and it only depends on the ECM stiffness and $k_l$. Panel B shows the H-VM simulations assuming a constant value of $k_l$ (elastic behavior) And for HepG2a (panel C), a non-linear strain behavior of the nucleus is observed, up to 120 hours of culture. It can be seen that after 72 hours the magnitude of the deformation of the nucleus of the primary cells is at least twice that of the nuclei at 36h. In the panels B and C it is possible to observe that by only using the active response of NE and a long.term elastic approach, the model does not reproduce the experimental strain behavior in time.

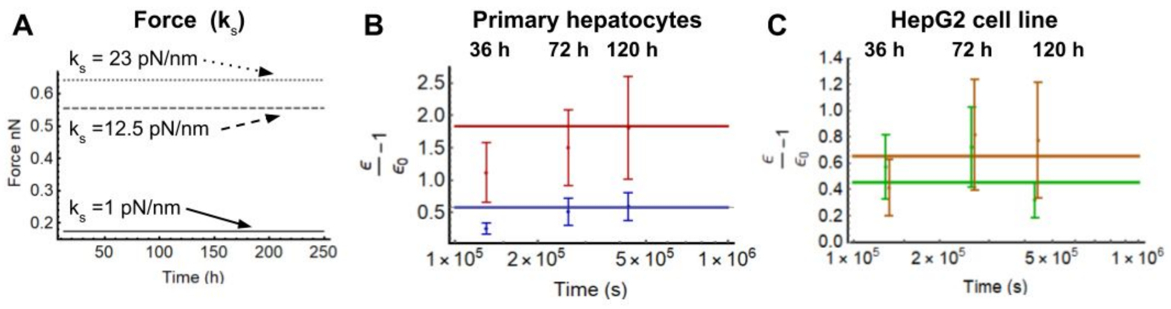


**Figure S1**. Long-term behavior vs elastic model.

A) Force as a function of time for three different physiologically-relevant substrate stiffnesses. In B) Strain of the nucleus of the primary hepatocytes against H-VM, assuming a constant $k_l$ behavior (long-term elastic behavior) on soft (blue) and stiff (red) substrates, which causes a constant deformation over time. C) Same graph for HepG2 cells on soft (green) and stiff (orange) substrates.

#### S.2 Viscoelasticity vs Hyperelasticity

In order to comprehend the limitations of the 1D model for a more scrupulous interpretation of the results, a finite element analysis (FEA) has been performed using the Structural Mechanics module of COMSOL Multiphysics version 5.5. The compression of a solid sphere with a 5 μm radius representing the cellular nucleus was simulated. The mechanical properties of the modeled sphere were parameterized to match the one of our proposed model at short timescales (see Figure 1B in the main text). Considering the reversible[9] and irreversible[10,11] behavior of the nucleus, a hyperelastic (Neo-Hook) model was also simulated with FEA. Figure S2A depicts the simulated setup for the nucleus of an hepatocyte compressed 20% in the *z* (vertical) direction, along the nucleus height (because, the Neo-Hookean model is optimal for the small strain regimen), with the following configuration. First, the sphere representing the cell nucleus was placed on top of a linear elastic substrate with a Young's modulus of 23 kPa and was compressed by a disc, which mechanical properties were chosen to resemble that of a silicon glass. Then, it was supposed that the contribution of osmotic stress remained constant over time; a valid hypothesis as it is known that osmotic stresses contribute to the resistance of the nuclear deformation due to compression[12]. Also, the influence of nuclear pores was considered constant and linear with respect to the total deformation and may then be represented by a scale factor $\varphi$ (Equation 3 in the main text).

In Figures S2B and S2C the distributions of stresses over the complete nuclear shape are presented and compared for an hyperelastic and viscoelastic material. As expected, the two behaviors are very different. The order of magnitude of the stresses remained constant in time for the nucleus that was modeled as a hyperelastic material, which is contrasted with the magnitude of stress at t = 0 s and the decrease at t = 45 s in case of a viscoelastic model of the same sphere. Moreover, it is observed that in both cases there is a clear transmission of stress to the substrate on which the nucleus was placed and compressed. The results also suggest that nuclear compression dissipates or stores elastic energy depending on the mechanical properties of the cell substrate. Also, it is found that the magnitude of the deformation is dependent on the mechanical properties of the substrate. Figure S2D presents the relaxation modulus on the transverse plane at half height for the FEA simulation and it is compared to the relaxation modulus calculated for the 1D model that is presented in Figure 1C. It may be observed that both 1D approximation and 3D FEA model predict an approximate value of 0.6 after 40 s of compression, representing a 40% relaxation of the stress exerted on the modeled nucleus. As no temporal shift was observed between 1D and 3D, it is appropriate to say that the 1D model may be employed to gain knowledge of the tendency of the mechanical behavior of the nucleus in 3D.

Finally, the simulations shown in Figure S2B and S2C demonstrate that one of the fundamental differences between the assumption of a dissipative versus a non-dissipative nucleus is that the stress suffered by the cell nucleus will diminish in minutes or seconds in spite of being under a constant deformation.

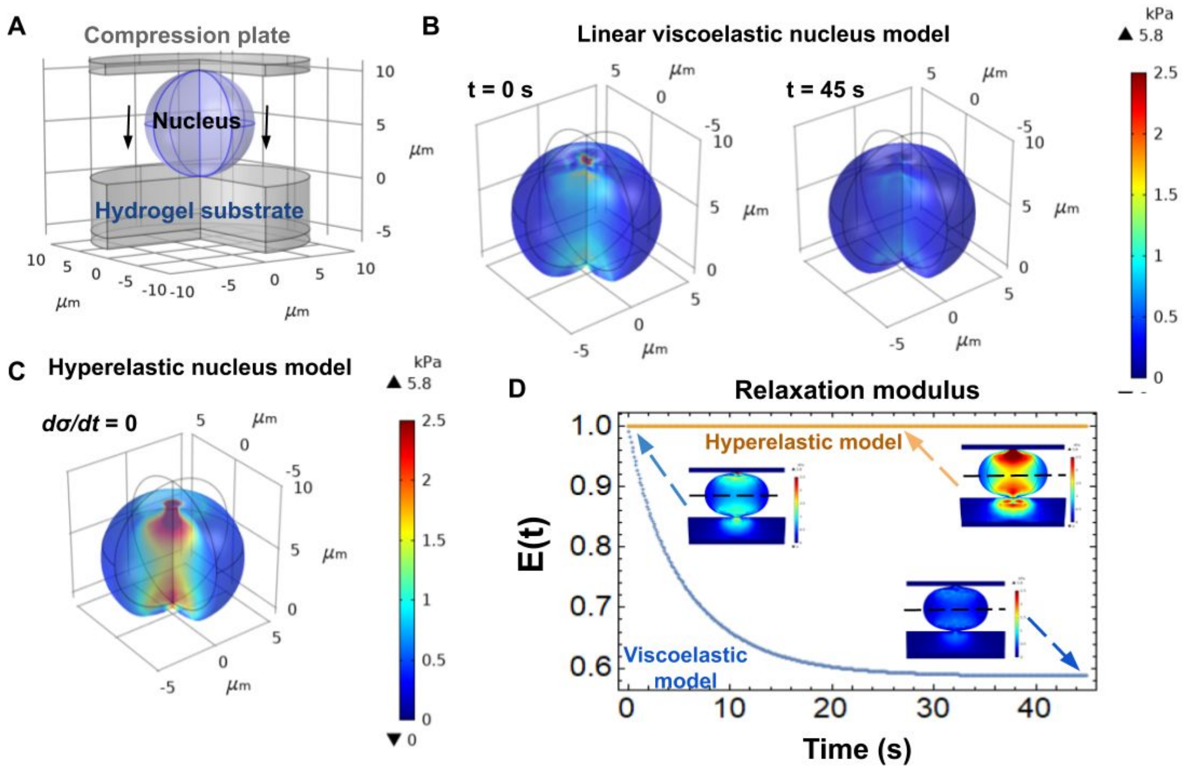


**Figure S2.** Impact of viscous dissipation on the mechanical characteristics of a 3D cell nucleus. Using a finite element model simulation performed with the Structural Mechanics Module of Comsol Multiphysics v5.5, the compression of a solid sphere was simulated. A linear viscoelastic (two-arm generalized Maxwell system) and a hyperelastic (neo Hook) model were used for comparison.
A) Setup used for compression analysis: cell nucleus between a linear elastic substrate with a 23 kPa Young's modulus (emulating a stiff hydrogel) and a very stiff glass disc. The disc is displaced down 2 µm vertically, compressing the nucleus 20%.
B) Evolution in time of the dissipation of the distributed stresses inside the nucleus at 0s and 45s, when the nucleus is simulated by a linear viscoelastic model.
C) Dissipation of the distributed stresses inside the nucleus when simulated as a hyperelastic material. As expected, there is no temporal dependency of stresses distributed inside the nucleus.
D) Relaxation modulus of the nucleus (on the transverse plane). The dissipative behavior of the nucleus not only decreases the stress upon it but also diminishes the stress transferred to the substrate.

#### S3. Program for the automation of characterization and segmentation of the nuclei

Experimentally, the affine deformation of a body contains all the information about its geometric and mechanical properties, regardless of the theoretical model used to interpret this information. In the context of cell biology, it is very common that the quantification of the certain parameters of the cell (or nucleus), such as its shape, projected area, height and aspect-ratio are carried out manually. This arduous work typically limits the number of objects characterized and can introduce some bias into the data. In the particular case of the nuclei of liver cell used here, it was necessary to sample a significant number of nuclei of each condition to characterize, since there is a great heterogeneity in the sizes and shapes. This is why an algorithm was programmed in Python 3 for the automated segmentation of stacks, with a good control for the operator. More importantly, this program aims at limiting the possible introduction of bias by the operator during the characterization of data.

*Confocal microscopy image processing (segmentation and quantification of fluorescence intensity)*: for image analysis, 3 to 5 images were used per sample and all the conditions were performed in duplicates (at least 6-10 images per condition). The images were analyzed in a Google Colab notebook using Python 3 language (programs are available upon request). Nuclei were segmented following the following procedure: each image stack was splitted in its three component channels (denoted as R, G, and B for red, green and blue respectively). From the R and G stacks, two new stacks were obtained, denoted as R' and G', through the moving average technique[13], with which averages of subsets are computed from an observations set. In this case, the observations consist of each of the images conforming the stack. In the next step, a new S stack was obtained through the computation of S = R'- G', and to each image in S a Gaussian filter was applied in order to reduce the noise, followed by an Otsu thresholding; both of them were implemented through the GaussianBlur and threshold functions from OpenCV library[14], giving as a result a new stack formed by binary images where white pixels conformed the objects, some of which related to nuclei positions. Because some nuclei may be in contact with others, thus generating only a unique object in the binary image, a watershed algorithm [15] was used in each binary image in the stack in order to separate the objects: the result was a new stack formed by images with labeled pixels, where each label indicates belonging to an specific object in the binary image.

After obtaining the binary stack, three filters were applied to each image in the stack, in order to retain only the objects related to the nuclei: a number-of-objects filter, an area filter and a circularity filter. The first filter allows to analyze only the properly thresholded images. The second filter computes the area $A$ of the objects, in squared pixels, through the function contourArea from the OpenCV library, then this value is converted to the image scale. Because the nuclei areas belong to a defined range, only the objects falling in this interval are considered (20 - 500 um). The last filter computes the circularity $C$ of the objects through the equation $C = 4\pi\frac{A}{P^2}$ [16], where $P$ is the perimeter of the object in pixels and is computed with the arcLength function from the OpenCV library. Also, this value was converted to the image scale. Circularity value reach a maximum value of 1 if the object is a circle, an a value of $\pi/4$ if the object is a square [17]. For this reason the objects with $C > 0.8$ where considered as related to some nuclei, although in some cases it was necessary to take into account lower values, with a minimum of $C = 0.65$.

In addition to the computed shape descriptors used in the filters (area, perimeter and circularity), the eccentricity $ecc$ was computed from the fitted ellipses to the objects that passed the three filters. This ellipses were obtained through the fitEllipse function from the OpenCV library, and using the equation $ecc = \sqrt{1-(b/a)^2}$, where $a$ and $b$ are the major and minor axis of the fitted ellipse.

Moreover, from each object that passed the three filters the centroid was computed using statistical moments[18]. Having all the centroids from all the binary images in the stack, the program ends up with a set of points in a 2D space where clusters are observed; those clusters indicate the belonging of several objects to one nucleus. Cluster classification is made with a mean shift algorithm[19,20], using the MeanShift function from the scikit-learn library[19,20]. This classification assigns a set of objects to an unique nucleus and at the same time finds the centroid of the objects set.

Regarding the quantification of fluorescence intensity, a projection in the Z axis was performed for every set of objects belonging to the same nucleus, obtaining as a result a new object which was fitted to a circle using the minEnclosingCircle function from the OpenCV library, hence obtaining values of a radius $r$ and a center $c$. Based on the $r$ value, two circles were generated with radius of $r' = 0.8r$ and $r'' = 1.2r$, and a center equal to the centroid found with the mean shift algorithm. The mean intensity within both circles is then computed for the images in the R stack, something similar was performed for the G stack but in this case the radius of the circles was fixed to 12 and 20 µm, using the same centers as in the R stack. Those values were used because 12 µm is the average radius of a primary hepatocyte in 1kPa after 2 hours of culture (this value was calculated with the experimental information in this article), and 20 µm is equivalent to 4 times this area. The use of two different radii was implemented in order to analyze if the IF trend depends on the area where it is quantified.

The validation of this procedure is finally achieved by comparing the areas computed by the program against areas quantified using ImageJ by manual delimitation of at least 200 nuclei per condition (Supplementary material S7). Interestingly, as a validation of our implemented program, it was observed that data obtained from manual analysis (approximately 200 nuclei) consisted of a subset of all the nuclei quantified by the automated program (approximately 800 per condition).

The outputs of the program are:

- an image with the z-projection of each stack,
- an image with the contours segmented in each z-plane and the labels associated with each segmented nucleus (these labels had the purpose of maintaining traceability, since it is possible to trace each specific nucleus that was characterized, for further review after analysis),
- two images with the areas in which the fluorescence intensity of the DAPI channel and the CK-18 channel were measured
- a *CSV* file that contains the tag data, the z position, area, perimeter and shape factor or circularity of each segmented contour.

Figure S3 summarizes all the steps used to segment the liver cell nuclei and obtain the *CSV* file. From this segmentation, the circularity (or shape factor) the projected area and the fluorescence intensity were measured. This program is shared upon request. NOTE FOR REVIEWERS: IT WILL BE SHARED IN A GITHUB REPOSITORY UPON PUBLICATION OF THIS PAPER.

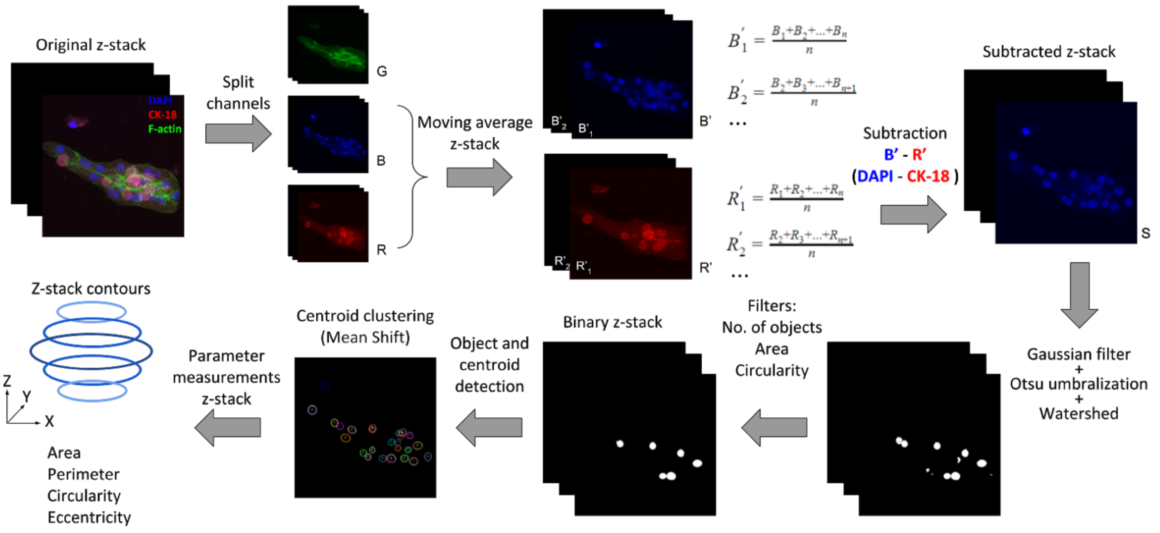


**Figure S3.** Steps of analysis for nucleus segmentation from confocal images.
After the original z-stack is splitted in its three RGB color components, the moving average of the B and R channels, corresponding to DAPI and CK-18 stains, are computed, generating the new stacks R' and B'. Next an element wise subtraction B'-R' was calculated, where each element is an image in each stack, creating a new z-stack denoted by S. Afterwards, to every image in the stack S a Gaussian filter was applied, followed by an Otsu thresholding and a watershed algorithm, giving as a result a new stack formed by binary images that contain the nucleus and some residual objects from the thresholding. This particular (noise) objects were removed with the help of three filters, the first one avoids to revise binary images with numerous objects (usually a result of a bad thresholding), the second filter removes objects that are too small or too big for the size of a nucleus and the third filter removes objects which are too far from a circular shape. This results in a new z-stack conformed by binary images containing the objects to analyze. In order to classify all the objects belonging to one nucleus, every centroid of the objects is computed and all of them are projected in a 2D plane where a mean shift algorithm is used to perform a centroid clustering, allowing to label the objects to their corresponding nuclei. Finally, for every contour object, the corresponding area, perimeter, circularity and eccentricity values were computed and saved in a file for later use, listing all processed nuclei for further review of the analysis.

#### Image processing of fluorescence images

For this procedure from 6 and up to 10 images were used per condition and all the conditions were done in duplicates. As the confocal images, these images were analyzed in a Google Colaboratory notebook using Python 3 language. Nuclei were segmented under a procedure similar to the used for the confocal images. In the first step the images were splitted in their three channels, keeping only the blue channel. The next step is to apply a Gaussian filter to the image to blur it, followed by a weighted adaptive threshold using a Gaussian window. This was implemented through the adaptiveThreshold function from the OpenCV library, the output was a binary image with white objects over a dark background. Afterwards, morphological operations of opening and closing were applied to the binary image, opening operation removes small objects while closing operation removes holes within the objects. The resultant binary image then passes through two filters, one related to the area and one related to the circularity, both similar to the filters used in the processing of the confocal images and mentioned above.

This procedure was applied to the image twice, using different window sizes for the Gaussian filter and different values for the area filter. This was done in order to adapt the program adequately to the different nuclei sizes. Computation of the area, perimeter, circularity and eccentricity was done over the objects that remained in the last binary image, similar to the processing of the confocal images. The data generated from this program was also exported in “csv” files and for every image analyzed one image is created showing the labels and contours of every segmented nucleus in the image.

All the *CSV* files were finally summarized and analyzed using Wolfram Mathematica 12.1. For each condition, a histogram was generated together with its probability density function (PDF). If the adjusted PDF consisted of a mixed distribution, then a deconvolution was performed to find the characteristic sizes present in each condition. Given the nature of the phenomenon studied here, we only fitted continuous PDFs (Normal, LogNormal, Logistics, Gamma, Extreme-Value and Mixture Distributions). Figure S4, S5, S6, S7 and S8 show a summary of the complete characterization of the area and height of the cell nuclei studied here in each condition, together with the shape factor and the nuclear aspect ratio. The aspect ratio was calculated in two ways: (a) as the ratio between the maximum height and the maximum length crossing the projected area or (b) as the ratio between the maximum height and the square root of the projected area. AS can be observed in Figure S8, the only difference between both methods lies in the magnitude (the value) but their time behaviors are similar.
Regarding the results obtained for the aspect ratio (Figure S8 A and B), the observed behavior indicates a volume decrease for primary cells on both stiffnesses and for HepG2 on the stiff substrates (the volume increases after 72h in this case).

A comparison between the results obtained rapidly and in high volume with our developed program and results obtained manually (in lower volume, by an operator, using ImageJ software) was also performed and is presented in Figure S9 for the analysis of at least 100 nuclei of primary hepatocytes. Panel A shows that the same trends as those measured using the automatic segmentation program were found. In panel B, an increase of the nucleus size of at least 100% can be observed in the rigid condition at 36 h and it increases with time. In panels C) and D) it is shown how the nuclei tend to a more ellipsoid shape in time in the rigid condition, while in the soft condition they remain almost constant. This also correlates with the results obtained with the automated program. Finally panel E shows a comparison between the projected area obtained with the two methods. As can be observed, there is no significant difference between them and this thus validates the automated quantification procedure (for a much larger volume of nuclei in less time). In order to find out if the difference was significant in panel A and E a one-way Ordinare ANOVA test with Tukey’s correction for multiple comparison was performed and a $p<0.05$ value was considered as significant.

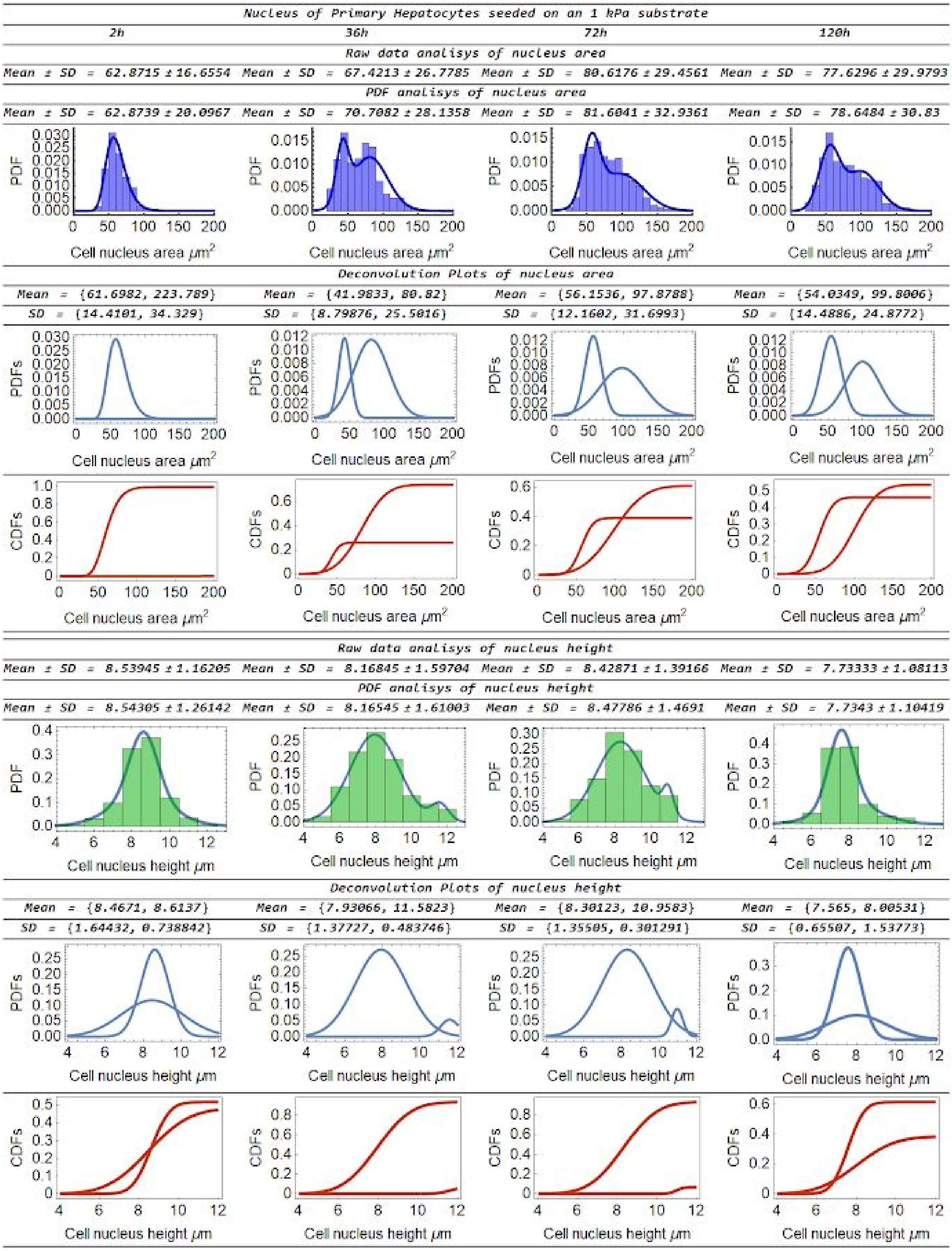


**Figure S4.** Quantification and deconvolution of the projected area and height of primary hepatocytes on soft hydrogels fixed at 2, 36, 72 and 120 hours.

The height of each bar in the histogram is the value of the PDF adjusted to each condition. The width was taken at 10µm for the area and 1µm for the height. The CDF value associated with each deconvolution determines the dynamics between populations. In this case, the dominant population is the one suffering a greater change in its projected area.

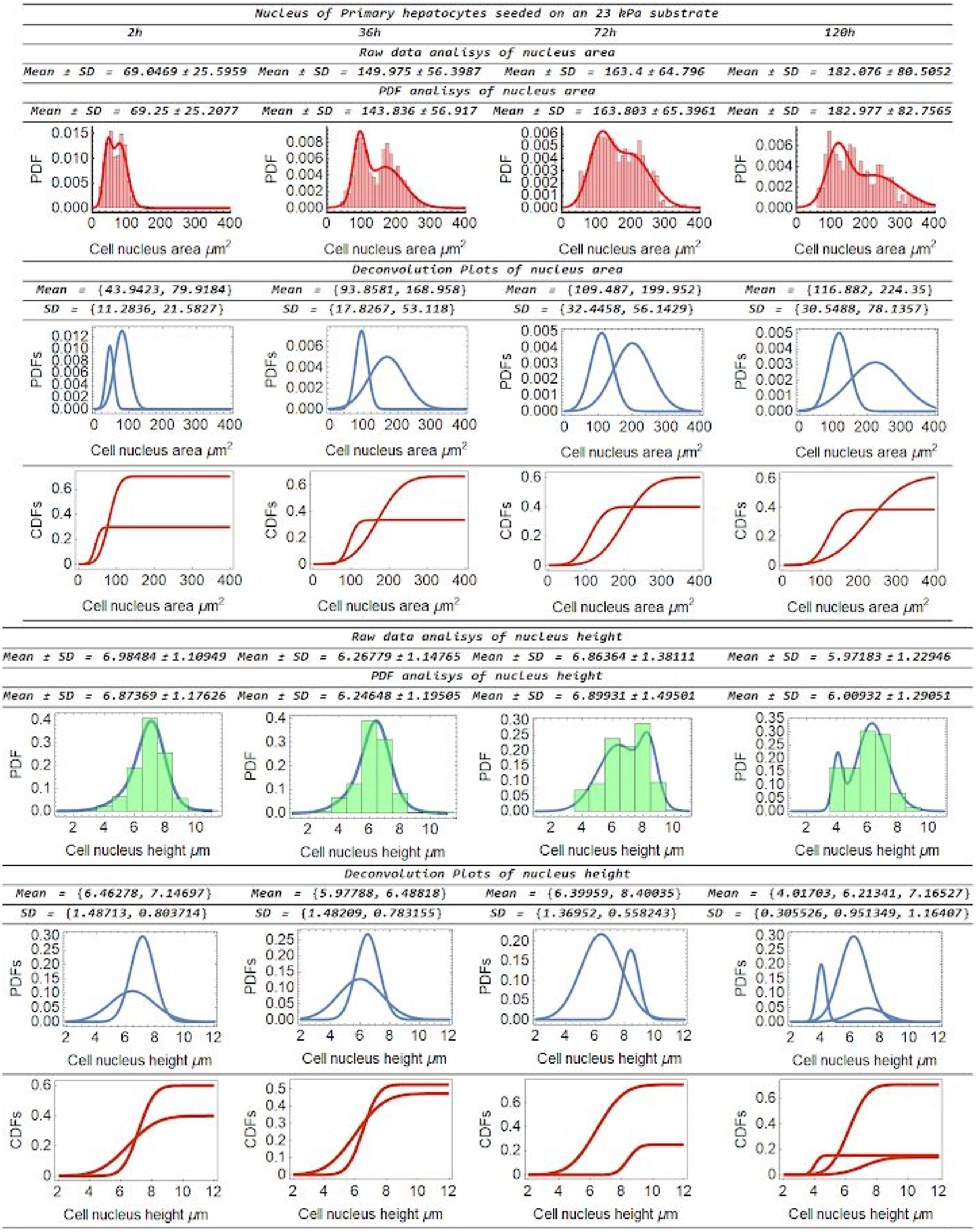


**Figure S5.** Same results than Figure S4 but for a stiff (23 kPa) substrate.

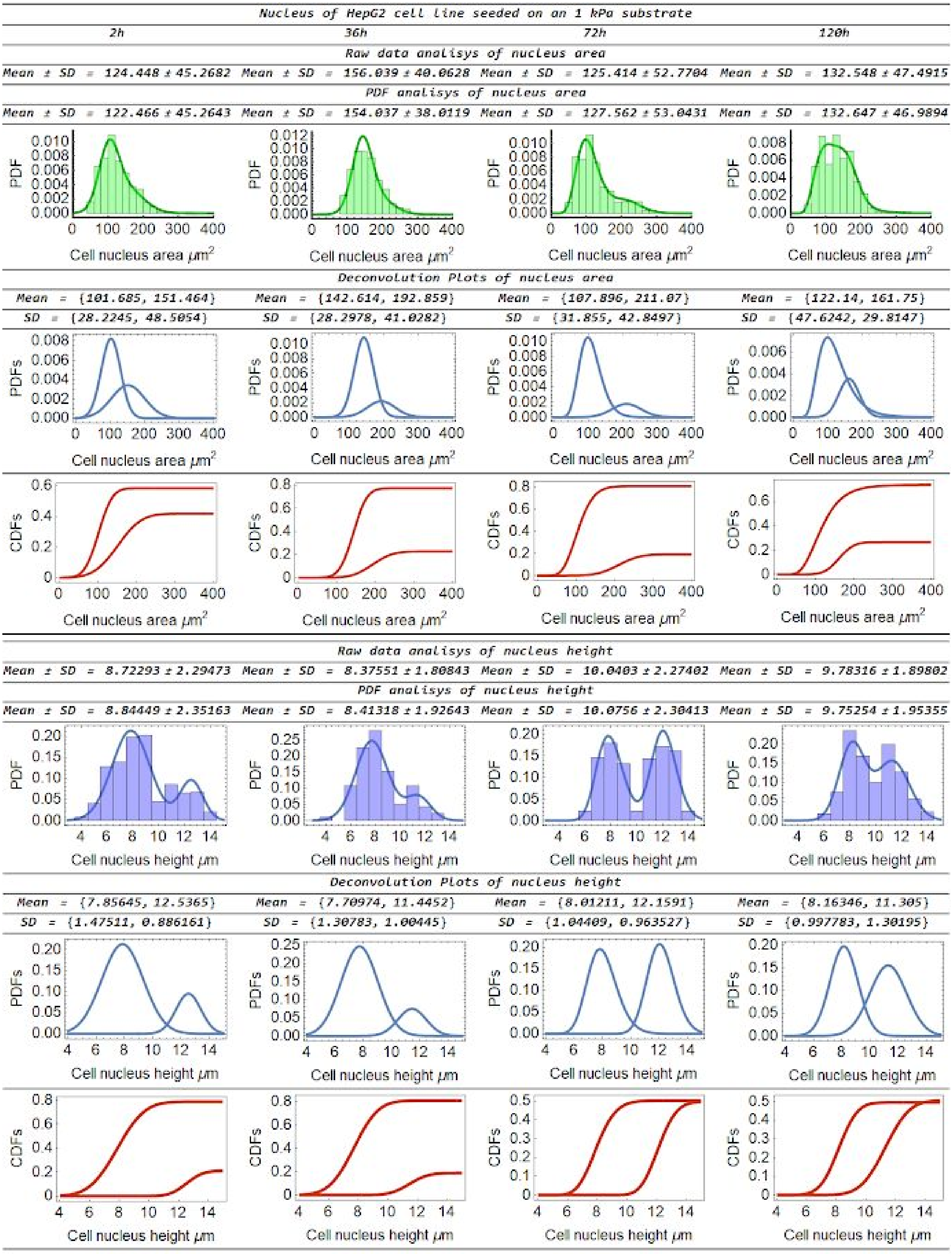


**Figure S6.** Same results than Figure S4 but for HepG2 cells. The width of each bar was taken at 15μm for the area and 1μm for the height. The CDF value associated with each deconvolution determines the dynamics between populations and in this case, the number of nuclei incrementing their projected area increases at 72h and decreases again at 120h, being as probable as the population that did not modify its area. This temporal behavior is also observed for nuclei incrementing their heights in time.

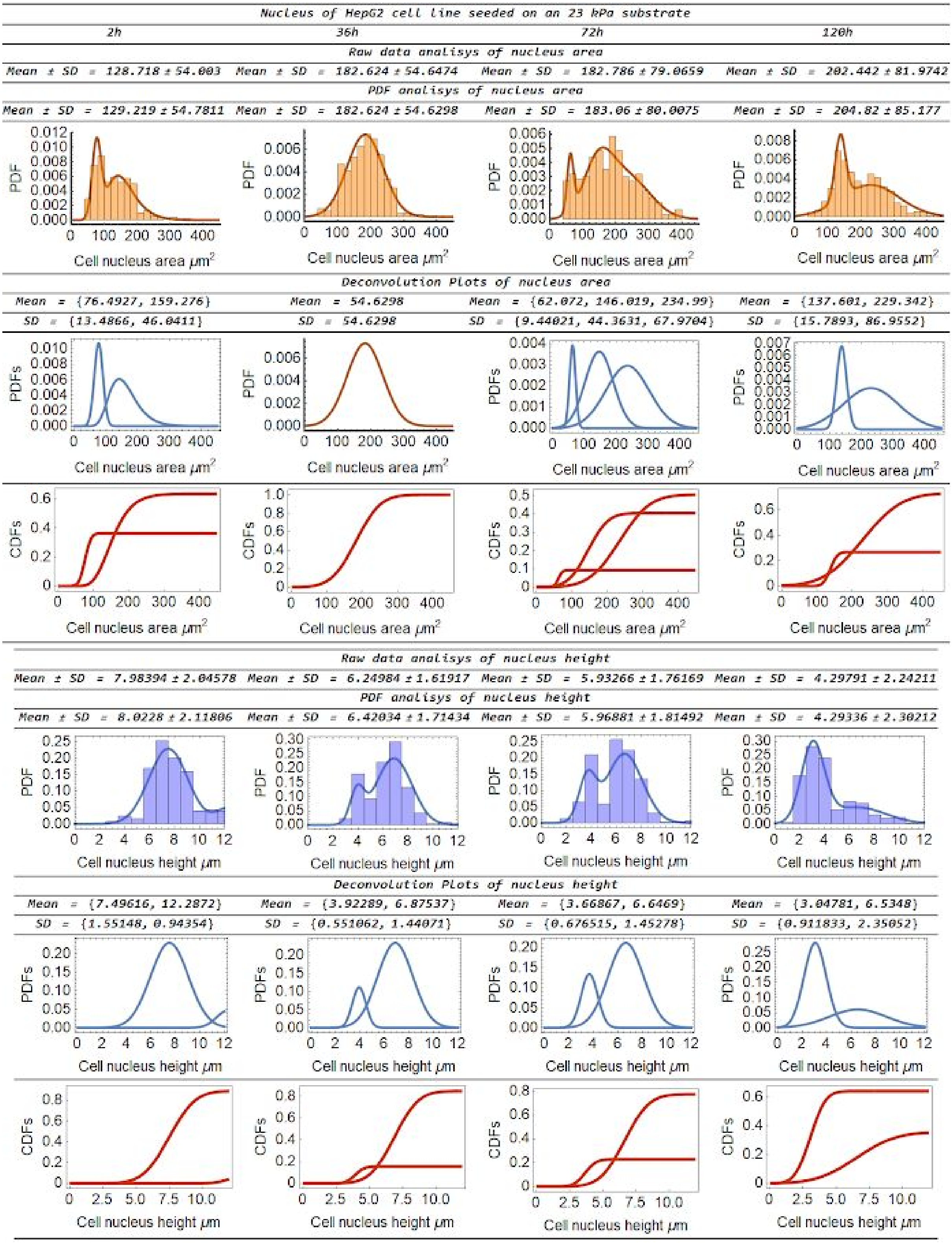


**Figure S7** Same results than Figure S6 but for a stiff substrate. In this case, All cell populations incremented their projected area until 72h and decreased it at 120h. However, contrary to the soft condition, here the nuclei decrease their heights.

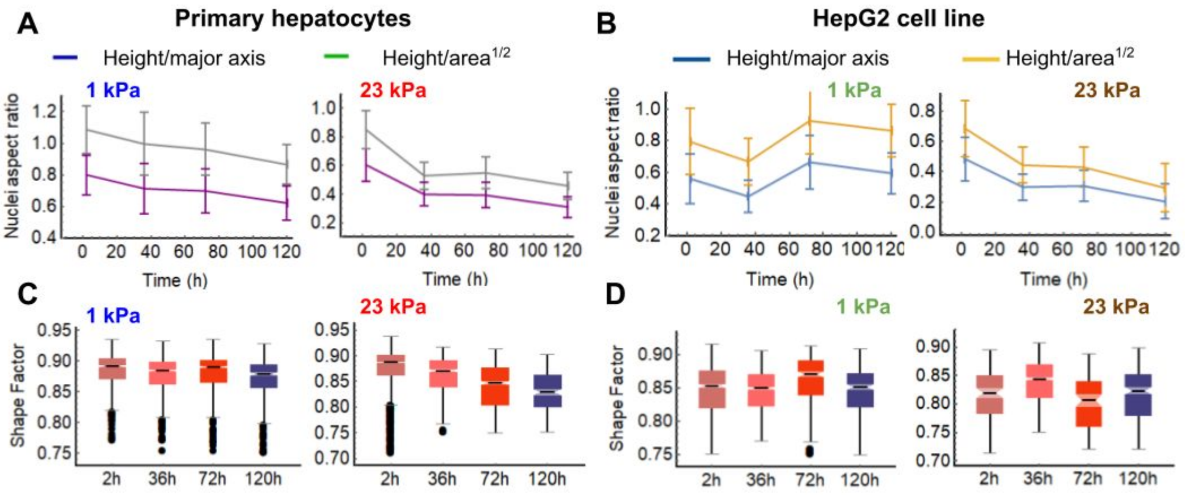


**Figure S8**. Morphology characterization of hepatic cells in culture.

A) Nuclear aspect ratio (AR) for primary hepatocytes seeded on soft (1 kPa) and stiff (23 kPa) hydrogels. The major AR change is seen for hepatocytes seeded of a stiff substrate. B) Nuclear AR of HepG2 cells. In this case, on soft substrates it decreases before 72 h of culture and increases again (reflected in the decrease of the Poisson's ratio) and on stiff substrates it keeps decreasing until the end of the experiment. C) and D) Nuclear shape factor $sf = 4\pi P/\sqrt{A}$ (*P* is the perimeter and *A* the area of the nucleus) of hepatocytes and HepG2 cells on soft and stiff substrates. These results suggest a decrease in nucleus circularity of both cells when the stiffness increases.

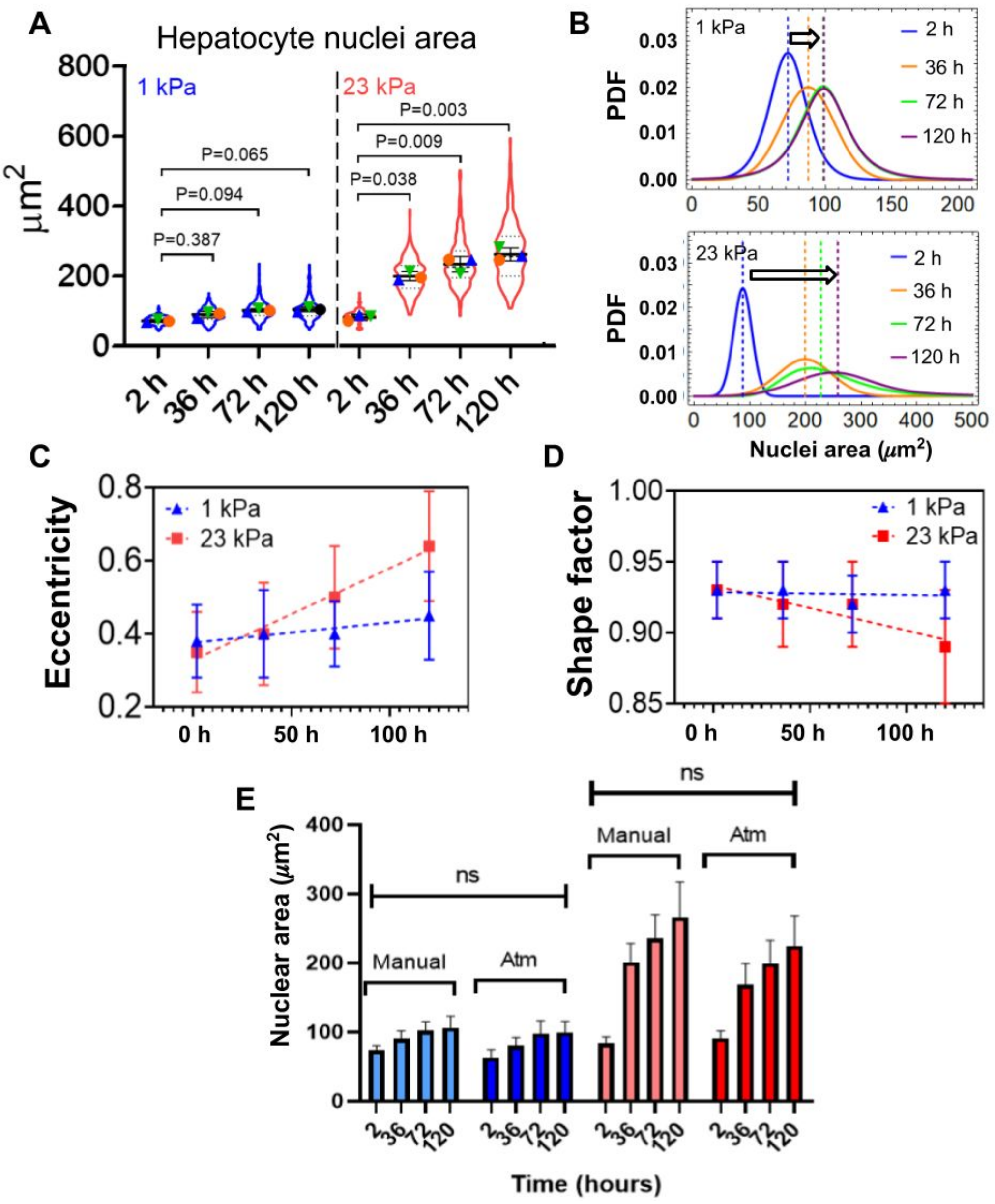


**Figure S9** Manual characterization of primary hepatocytes nuclei. Using the software ImageJ the contour of the projected area was delimited manually. A) Projected area of the nuclei in the different stiffness conditions, at least 100 nuclei were quantified per condition. B) Comparison of the PDF of the soft and stiff conditions over time. C) Evolution of the eccentricity of the nuclei for both conditions. D) Evolution of the shape factor of the nuclei for both conditions. E) Comparison of the projected areas obtained with the data quantified by hand (M) and the data quantified automatically (Py).

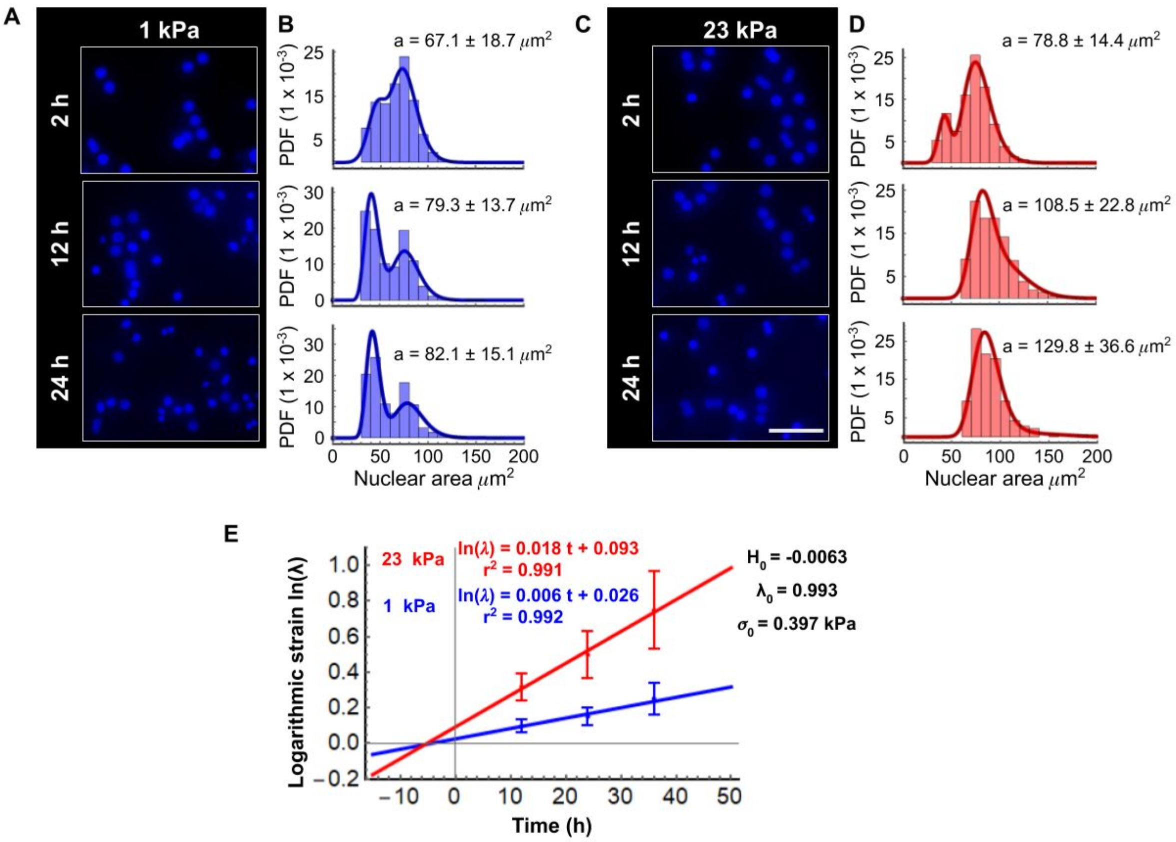


**Figure S10**. Primary hepatocytes at 2, 12 and 24 hours in culture. A) and C) are representative fluorescence microscopy micrographs of primary hepatocytes at 2, 12 and 24 hours in culture on polyacrylamide hydrogels of 1 kPa and 23 kPa coated with type I collagen. B) and D) present the quantification of the projected area of the nuclei in the different stiffness conditions. E) Pre-strain in nuclear primary hepatocytes. Using Equation S13 for 12, 24 y 36 h the nuclear pre-strain was calculated and then the pre-stress was obtained knowing the value of $k_{l0}$ . Scale bars: 50µm.

Regarding the quantification of the intensity of fluorescence of CK-18, several distributions were used to identify the best fit. In both cases of primary hepatocytes and HepG2, it was found that the log-Normal and Inverse-Gaussian were better fits for the obtained histograms (Figure S10.1 and 10.2). It is possible to observe that the IF(CK-18) decreases on both stiffness conditions for primary cells, similar to the behavior of $k_l$. This can be associated with the volume change observed in the results of Figure S8. For HepG2, there is a distinction between soft and stiff conditions that may be associated with the dynamics of the Poisson's ratio found in Figures 5 and 6 of the main text and the AR results seen in Figure S8.

*Fluorescence intensity of ck-18 of Primary hepatocytes seeded on an 1 kPa substrate*

*Raw data analisys (r=12 μm)*

| | | | |
|---|---|---|---|
| Mean ± SD = 22.0081 ± 11.808 | Mean ± SD = 24.8402 ± 14.763 | Mean ± SD = 15.2499 ± 8.83532 | Mean ± SD = 16.3807 ± 9.76809 |

*PDF analisys (r=12 μm)*

| | | | |
|---|---|---|---|
| Mean ± SD = 22.3134 ± 11.9073 | Mean ± SD = 24.5513 ± 14.4632 | Mean ± SD = 15.0929 ± 8.7099 | Mean ± SD = 15.9165 ± 9.80949 |
| Mode = 26.154 | Mode = 20.807 | Mode = 12.4017 | Mode = 11.5017 |

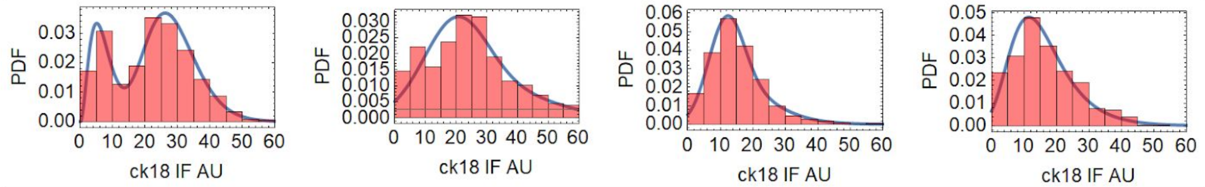


*Raw data analisys of (r=20 μm)*

| | | | |
|---|---|---|---|
| Mean ± SD = 15.4027 ± 8.26207 | Mean ± SD = 20.0484 ± 11.3806 | Mean ± SD = 13.4265 ± 7.34988 | Mean ± SD = 15.1738 ± 8.53483 |

*PDF analisys (r=20 μm)*

| | | | |
|---|---|---|---|
| Mean ± SD = 15.2964 ± 8.26681 | Mean ± SD = 21.1497 ± 13.5281 | Mean ± SD = 13.5757 ± 6.61523 | Mean ± SD = 15.3176 ± 8.51942 |
| Mode = 11.7985 | Mode = 12.327 | Mode = 10.3522 | Mode = 10.5792 |

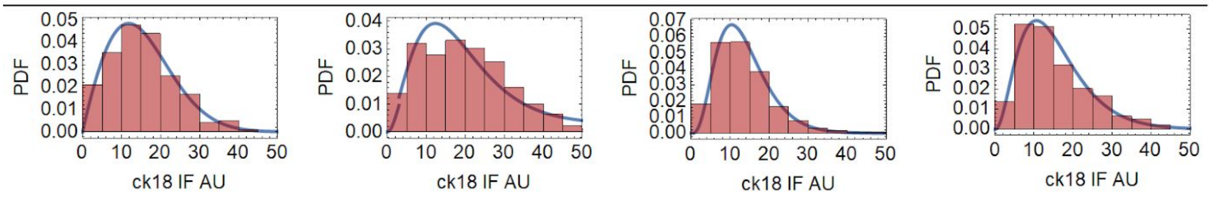


*Fluorescence intensity of ck-18 of Primary hepatocytes seeded on an 23 kPa substrate*

*Raw data analisys (r=12 μm)*

| | | | |
|---|---|---|---|
| Mean ± SD = 19.4628 ± 10.2089 | Mean ± SD = 10.1123 ± 7.442 | Mean ± SD = 10.3432 ± 7.06729 | Mean ± SD = 10.0489 ± 9.36527 |

*PDF analisys (r=12 μm)*

| | | | |
|---|---|---|---|
| Mean ± SD = 19.2071 ± 10.7475 | Mean ± SD = 10.2052 ± 6.94657 | Mean ± SD = 10.0597 ± 6.8638 | Mean ± SD = 10.1979 ± 9.62571 |
| Mode = 16.1405 | Mode = 7.07888 | Mode = 4.15172 | Mode = 1.99199 |

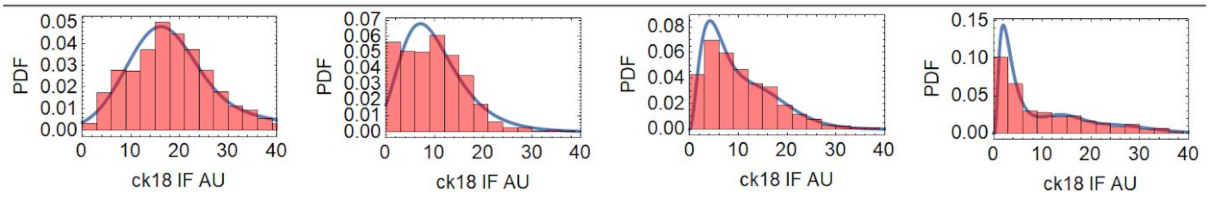


*Raw data analisys (r=20 μm)*

| | | | |
|---|---|---|---|
| Mean ± SD = 14.0229 ± 8.21371 | Mean ± SD = 9.58052 ± 6.41219 | Mean ± SD = 9.81099 ± 6.35621 | Mean ± SD = 8.89525 ± 7.23941 |

*PDF analisys (r=20 μm)*

| | | | |
|---|---|---|---|
| Mean ± SD = 13.7568 ± 7.97473 | Mean ± SD = 9.96899 ± 7.35646 | Mean ± SD = 9.81842 ± 6.34824 | Mean ± SD = 9.00126 ± 7.45157 |
| Mode = 9.13385 | Mode = 4.84551 | Mode = 6.96137 | Mode = 2.83257 |

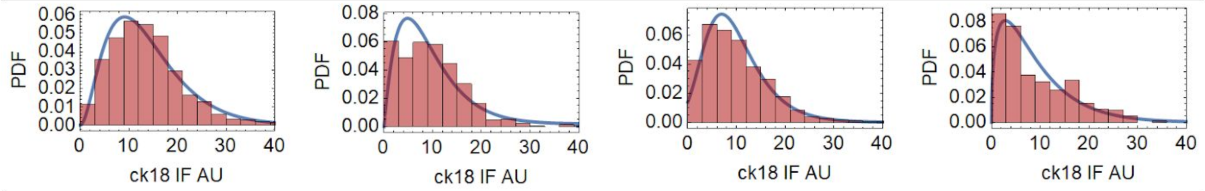


**Figure 11.1**. Quantification of the intensity of fluorescence of CK-18, IF(CK18), for primary hepatocytes.

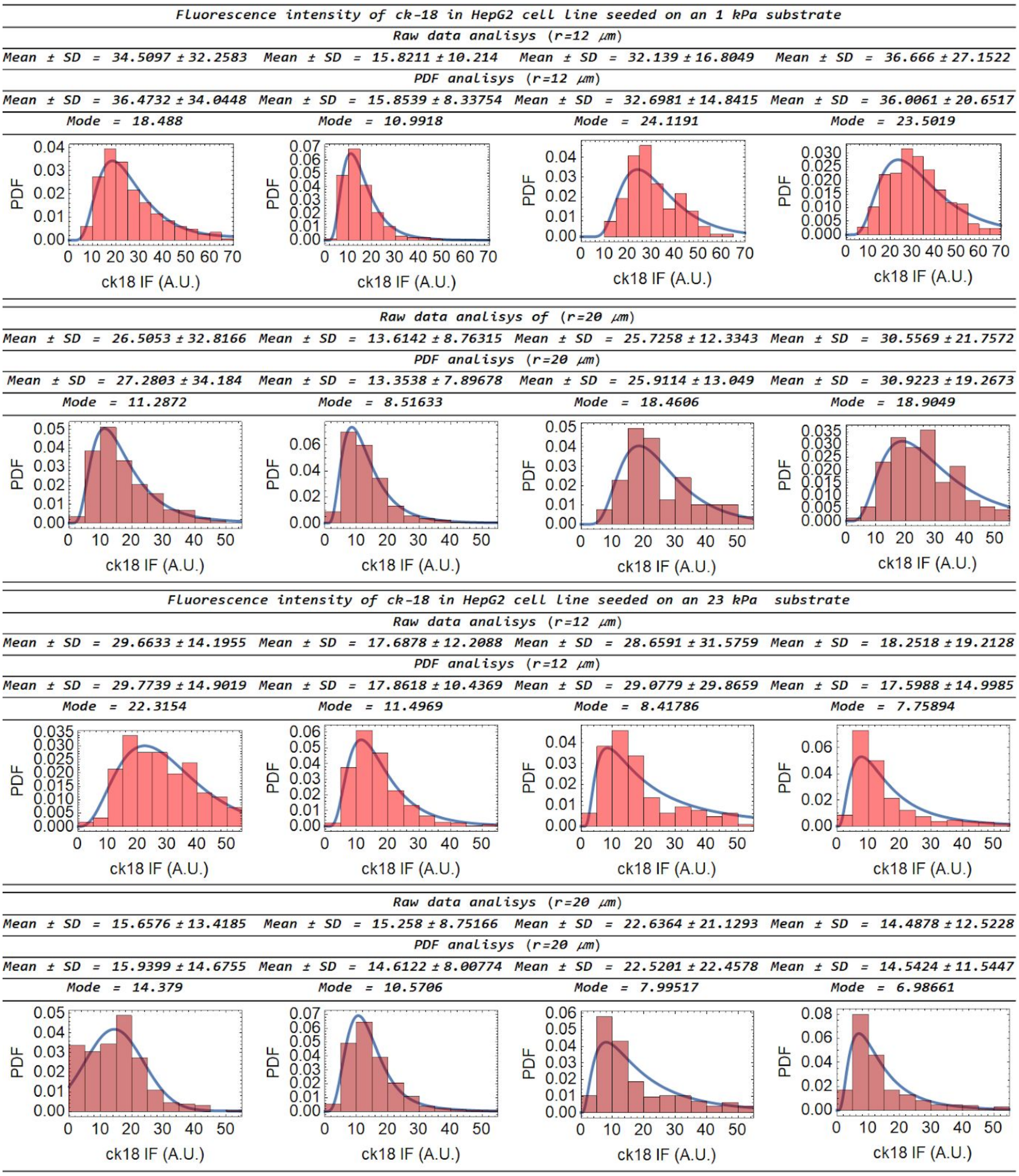


**Figure 11.2**. Quantification of IF(CK18) for the nuclei of HepG2 cells.

#### S3. Parameters Tables

##### Supporting Information Table S1

Stiffness and relaxation time of the several liver mechanical characterizations.

| Viscoelastic properties of some tissues and cells | | | |
|---|---|---|---|
| **Tissue** | $\tau_{NE}$ * (s) | **Stiffness E (kPa)** | **References** |
| Liver ECM | 0.05 | ~ 0.15 ± 0.08 | [21] |
| Liver tissue | 1.19 | 1.12 ± 0.15 | [22] |

| | | | |
|---|---|---|---|
| Liver cell | 0.64 - 1 | ~0.75 - 1 | [23] |
| Liver F cutoff | ~30 - 48 | ~9 - 12 | [24] |
| Liver F1 | ~27 | 8.3 | [25] |
| Liver F4 | ~129 | 23 (Range 4.5 - 50) | [25] |
| Normal lung | 18.20 | 6.5 ± 1.5 kPa | [26] |

*These values were calculated using the relation $\tau \sim [\text{Lamin A:B}]^{2.5}$ as mentioned in [27] and in this case we used the following: [Lamin A] ~ $E^{0.7}$ [Lamina B] ~ $E^{0.2}$ where E is the value of the micro-stiffness. The direct relation is $\tau_{NE} \approx E^{1.75}/E^{0.2}$ .

**Supporting Information Table S2**

Basal parameters for the cell mechanics simulation

| Focal adhesion elements | | | |
|---|---|---|---|
| **Parameter** | **Description** | **Value** | **Ref** |
| $k_p$ | Plaque stiffness | ~1 pN/nm | [28] |
| $k_c$ | Effective spring constant of the clutch (integrin stiffness) | ~5 pN/nm | [29] |
| $k_s$ | Substrate stiffness: typical micro-stiffness of the tissues | ~0.2-40 pN/nm | [27] |
| $d_c$ | Integrin spacing | ~100 nm | [28] |
| $L$ | FA length | ~1000 nm | [30] |
| **Nucleus elements **** | | | |
| $k_{\mu T}$ | Stiffness of the microtubule | ~1 pN/nm | [1,31] |
| $k_a$ | Stiffness of the F-actin | ~1 pN/nm | [32–34] |
| $\rho_0$ | Initial myosin motor density (prestress) | ~0.5 pN/nm | [35,36] |
| $\beta$ | Chemo-mechanical coupling parameters related to the molecular mechanisms that regulate the engagement of motors | ~2.5 | [37–3929,38,40] |
| $\alpha$ | Chemo-mechanical coupling parameters related to the molecular mechanisms that regulate stress-dependent signaling pathways | ~1.5 | [29,40,41] |
| **Nucleus elements **** | | | |
| $k_{l0}$ | Nuclear stiffness (elastic behavior) | 2-25 pN/nm | [42] |
| $\varphi$ | Nuclear pore complexes fraction | ~0.2 | [43] |
| $k_{LB}$ | Stiffness associated with lamin B | 0 - 5 pN/nm | [27] |
| $\eta_{LA}$ | Viscosity associated with lamin A/C | 0 - 160 pN*s/nm | [27] |
| $k_{Ch}$ | Stiffness associated with chromatin & subnuclear elements | ~1 pN/nm | Estimated from [44] |
| $\eta_{Ch}$ | Viscosity associated with chromatin & subnuclear elements | ~5 pN*s/nm | Estimated from[45] |

* The estimations of the elastic modulus of chromatin presented here were calculated using a scale factor and the approach that uses Boussinesq-Green's function [5,6]. The scale factor used for the focal adhesions was 0.5 µm and 5 µm for the nucleus.

** These values were calculated using the relation $\tau$ ~ [Lamin A:B]$^{2.5}$ as mentioned in [27] and in this case we used the following: [Lamin A] ~ $E^{0.7}$ [Lamina B] ~ $E^{0.2}$ where E is the value of the micro-stiffness. The direct relation is $\tau_{NE} \approx E^{1.75}/E^{0.2}$ .

##### Supporting Information Table S3

Parameters fitted to equation 10.1 and 10.2

| | Primary hepatocytes | | | |
|---|---|---|---|---|
| | Correlation coefficient $R^2$ | | AIC* | |
| | Equation 10.1 | Equation 10.2 | Equation 10.1 | Equation 10.2 |
| **Soft (1 kPa)** | 0.999 | 0.999 | -4.77 | .-11.11 |
| **Stift (23 kPa)** | 0.999 | 0.993 | -3.21 | 12.37 |
| | **HepG2 cell line** | | | |
| | **Correlation coefficient $R^2$** | | **AIC*** | |
| | **Equation 10.1** | **Equation 10.2** | **Equation 10.1** | **Equation 10.2** |
| **Soft (1 kPa)** | 0.980 | 0.997 | 9.60 | 3.34 |
| **Stift (23 kPa)** | 0.989 | 0.999 | 17.05 | 1.43 |
| | **Primary hepatocytes** | | **HepG2 cell line** | |
| **Parameters Values** | | | | |
| **Parameter** | **Soft (1 kPa)** | **Stift (23 kPa)** | **Soft (1 kPa)** | **Stift (23 kPa)** |
| ***a*** | 0.202 | 0.067 | 0.565 | 0.053 |
| ***b*** | 6.5 x $10^{-7}$ | 3.1 x $10^{-7}$ | -5.5 x $10^{-7}$ | 1.2 x $10^{-7}$ |
| $x_{eff}$ | ~1.99 | ~1.99 | ~1.99 | ~1.99 |
| ***a'*** | 4.65 | 16.80 | 5 | 24 |
| ***b'*** | 1.03 x $10^{-5}$ | 6.31 x $10^{-5}$ | 2.51 x $10^{-5}$ | 8.70 x $10^{-5}$ |
| ***c'*** | 1.01 x $10^{-11}$ | 8.37 x $10^{-11}$ | 4.87 x $10^{-11}$ | 1.29 x $10^{-11}$ |

*The akaike information criterion (AIC) is used to estimate the quality of the fitted model[7].The model which AIC value is closest to 0 is better.
